# Novel Cln8 p.R24G mouse line replicates major clinical features of Northern epilepsy

**DOI:** 10.64898/2025.12.02.690107

**Authors:** J. Müller-Niva, M.H. Salo, S. Häkli, I. Gureviciene, A.-S. Makiou, H. Koivisto, H. Elamaa, I. Miinalainen, A. Heikkinen, L. Eklund, P. Sipilä, S. Kuure, H. Leinonen, J. Uusimaa, H. Tanila, R Hinttala

**Affiliations:** Research Unit of Clinical Medicine, Medical Research Center, University of Oulu, Oulu University Hospital, Finland; Biocenter Oulu, University of Oulu, Finland; A.I. Virtanen Institute for Molecular Sciences, University of Eastern Finland, Kuopio, Finland; School of Pharmacy, University of Eastern Finland, Kuopio, Finland; ECM-Hypoxia Research Unit, Faculty of Biochemistry and Molecular Medicine, University of Oulu, Finland; Institute of Biomedicine, Research Centre for Integrative Physiology and Pharmacology, Turku Center for Disease Modeling, University of Turku, Finland; Stem Cells and Metabolism Research Program Unit, Faculty of Medicine, University of Helsinki, Finland; GM-unit, Laboratory Animal Centre, University of Helsinki, Finland; Department of Children and Adolescents, Division of Paediatric Neurology, Oulu University Hospital, Oulu, Finland

**Keywords:** neuronal ceroid lipofuscinosis, neuroinflammation, neurodegeneration, epilepsy, disease model

## Abstract

**Rationale:** Northern epilepsy belongs to a group of genetically diverse lysosomal storage diseases, the neuronal ceroid lipofuscinoses (NCLs). A characteristic feature of NCL pathology is the accumulation of autofluorescent ceroid lipofuscin in the central nervous system. Northern epilepsy is a late-infantile-onset disease. Patients develop normally until 5–10 years old, when they first present with general tonic-clonic seizures, followed by progressive cognitive impairment and a decline in motor skills. Northern epilepsy is caused by a missense variant in *CLN8*, causing a p.R24G amino acid substitution. CLN8 deficiency has been studied traditionally using motor neuron degeneration (mnd) mice, which carry a spontaneous frame shift variant in the murine orthologue *Cln8*, which is not known to exist in humans.

**Methods:** We have generated the first Cln8 p.R24G mouse model (Cln8^R24G^) using CRISPR/Cas9. Phenotyping analysis of the mice was conducted using behavioral and histopathological studies focusing on the brain and retina.

**Results:** At birth, Cln8^R24G^ KI mice were viable and asymptomatic. As the mice aged, a progressive accumulation of autofluorescent ceroid lipofuscin containing the mitochondrial ATP synthase subunit C was evident in different brain regions and retinal layers. Health monitoring revealed that mutant mice developed progressive but mild motor symptoms around 7 months of age. Spontaneous epileptic seizures, like those observed in Northern epilepsy patients, were detected and recorded. An increase in FosB staining intensity, reflecting neuronal hyperactivity, was observed in hippocampal CA1-CA3 pyramidal and dentate granule cells and correlated well with the intensity of seizure activity. Neuroinflammation was evident at 4 months and increased dramatically with age, mainly in the thalamic VPN/VPL nuclei and moderately in the cortex. Neurodegeneration was most prominent in the VPN/VPL thalamic nuclei.

**Conclusions:** We have generated the Cln8^R24G^ mouse model that genocopies, for the first time, the pathogenic CLN8 variant present in patients with Northern epilepsy. These mice phenocopy major clinical features of the human disease, including mild motor impairment and, unlike the mnd mice, spontaneous generalized tonic-clonic seizures. We hypothesize that our mouse model will open new possibilities for developing and testing targeted treatment options for Northern epilepsy and, more broadly, for early-onset neurodegenerative disorders associated with epilepsy.

## Introduction

Northern epilepsy or epilepsy, progressive, with mental retardation (EPMR) (OMIM #610003) belongs to a group of neuronal ceroid lipofuscinoses (NCLs), also known as Batten disease. NCLs are a genetically and clinically heterogenous group of neurodegenerative disorders characterized by lysosomal accumulation of autofluorescent lipopigment storage material (Cárcel-Trullols et al., 2015; Haltia, 2003; Kousi et al., 2012). In general, the clinical course of NCLs includes progressive neurodegeneration, motor and cognitive decline, epilepsy, and loss of vision. Together, the NCLs are the most common cause of neurodegeneration in children (Ostergaard, 2016), with an incidence of 1 in 12, 500 live births in Scandinavia and an estimated incidence of 1 in 100, 000 live births worldwide (Santavuori, 1988).

Northern epilepsy is caused by a recessive variant c.C70G (rs104894064), leading to a p.R24G amino acid substitution in the CLN8 protein. The variant is enriched in the Finnish population, with a carrier frequency of 1 in 135 (Ranta et al., 1999). CLN8 encodes a transmembrane protein that localizes to the endoplasmic reticulum (ER) and ER-Golgi intermediate compartment (ERGIC). CLN8 has been described to participate in the transport of lysosomal enzymes from ER to Golgi via COP-coated vesicles (Bajaj et al., 2020; di Ronza et al., 2018; Lonka et al., 2000, 2004; Luzio, 2018). Other proposed functions include an involvement in lipid metabolism and homeostasis (Haddad et al., 2012; Passantino et al., 2013; Sheokand et al., 2025; Vance et al., 1997; Winter & Ponting, 2002) and cellular calcium homeostasis (Kolikova et al., 2011). Recently, CLN8 was shown to regulate lysosomal phospholipid composition by acting as a lysophospatidylglycerol acyltransferase involved in the biosynthesis of bis(monoacylglycero)phosphate or BMP (Sheokand et al., 2025).

Northern epilepsy is characterized by juvenile-onset epileptic seizures. The most common seizure type is generalized tonic-clonic seizures, though complex focal seizures and myoclonic seizures may also occur (Hirvasniemi et al., 1994). The frequency of seizures usually increases during puberty and decreases spontaneously in adulthood. Other symptoms of Northern epilepsy are progressive cognitive deterioration and motor impairment starting 2−5 years after seizure onset (Hirvasniemi et al., 1994, 1995). The disease progresses to intellectual disability by middle age. In contrast to other NCL forms, Northern epilepsy patients have a substantial life expectancy into their sixth or seventh decade (Haltia, 2003; Hirvasniemi et al., 1994).

Histopathological findings in autopsy samples from Northern epilepsy patients include a characteristic accumulation of autofluorescent, mitochondrial ATP synthase subunit C (SCMAS) immunoreactive storage material inside lysosomes, predominantly in neurons (Cárcel-Trullols et al., 2015; Haltia, 2003; Herva et al., 2000). Hippocampal areas CA2-4 are most affected (Herva et al., 2000; Tyynelä et al., 2004) as well as pyramidal cells in the cortical lamina III (Herva et al., 2000). An association between the accumulation of ceroid lipofuscin and neuronal loss, astrocytosis, and neurophagy have been documented in the CA2 area. Neuronal loss has also been detected in layer V of the isocortex. Astrocytosis and reactive gliosis indicating neuroinflammation have been modest in all brain areas except for the severely affected hippocampal area CA2(Herva et al., 2000; Tyynelä et al., 1993). Activation of microglia has been hypothesized to reflect the degree of neuronal loss in the hippocampus (Tyynelä et al., 2004).

More severe phenotypes associated with CLN8 variants were initially identified in Turkish families (Sahin et al., 2017). These patients suffered from epileptic seizures, motor and cognitive decline, and loss of vision. The disease progresses more rapidly than in Northern epilepsy patients. Other CLN8 variants associated with variable phenotypic disease features have also been described (Badura-Stronka et al., 2021).

Current preclinical knowledge on CLN8 disease progression has relied on the motor neurodegeneration (mnd) mouse model, which originated from a naturally occurring variant of *Cln8* (c.267-268 insertion C) resulting in a premature stop codon (Ranta et al., 1999). Mnd mice present with retinal degeneration (Bronson et al., 1993; Chang et al., 1994; Guarneri et al., 2004; Seigel et al., 2005), mild ataxia, and hindlimb clutching starting from 6 months of age, which progresses to severe spastic paralysis. Behavioral abnormalities include enhanced activity, aggressive features, and impaired habituation and memory starting from 4−5 months of age (Bolivar et al., 2002). The mice show brain atrophy and hydrocephalus (Pardo et al., 1994a) and suffer premature death at the median age of 10 months, with females perishing earlier than males (Holmes et al., 2022). Accumulation of autofluorescent storage material, SCMAS and lipids has been documented to occur in the neurons of mnd mice (Bronson et al., 1993; Chang et al., 1994; Cooper et al., 2015; Faust et al., 1994; Pardo et al., 1994), most prominently in the thalamic nuclei, hippocampus, piriform cortex, spinal cord and retina (Kuronen et al., 2012; Pardo et al., 1994). An increase in markers for microgliosis and astrocytosis has been evident in the brain and spinal cord in the pre-symptomatic phase preceding the onset of motor symptoms (Bronson et al., 1993; Fujita et al., 1998; Mennini et al., 2004). Gliosis and neuronal loss in brain follows a spatiotemporal pattern, and high vulnerability of the thalamocortical pathways, especially the somatosensory pathway (Kuronen et al., 2012b). In brief, the mnd mouse model phenocopies many features of Northern epilepsy, but it does not genocopy the causative pathogenic variant of patients. Further, while mnd mice are more susceptible to induced seizures (Kriscenski-Perry et al., 2013), spontaneous seizures − an important early symptom in Northern epilepsy patients − have not been reported to date.

In this paper, we describe generation and phenotypic characterization of the first *Cln8* knockin (KI) mouse model that genocopies the human Northern epilepsy variant, p.R24G.

## Materials and methods

### Mouse model

The Cln8^R24G^ mouse line was generated using CRISPR/Cas9-based genome editing (Cong et al., 2013; Mali et al., 2013). The crRNA used (ID:527230429) was selected with the CRISPR finder tool (Hodgkins et al., 2015). The ssODN (5’-ATGACTCCTGTGAGCAGCCACGGCTTGGCGGAGAGCATTTTTGACCTGGACTACGCTTCGTGGAAGATTGGGTCGACGCTAGCTGTTGCTGGCTTTGTCTTCTACCTGGGCGTCTTTGTAGTCTGCCATCAGCTCTCATCGTCCCTGAATGCCACCTA-3’, Ultramer^TM^ DNA Oligo, Integrated DNA Technologies, Coralville, IA, USA) contained CGG>GGG modification at codon 24 (ENSMUST00000027554, from the mouse genome assembly GRCm38) to create R>G change and silent mutations at codons 26 (ACT>AGC) and codon 28 (GCG>GCT) to create a *Nhe*I restriction site and to eliminate existing the PAM site, respectively. A preassembled Alt-R^TM^ crRNA/Alt-R^TM^ tracrRNA/Alt-R^TM^ S.p. HiFi Cas9 (Integrated DNA Technologies, Coralville, IA, USA) ribonucleoprotein complex, together with the ssODN, were injected (20ng/µl-20 ng/µl-20 ng/µl respectively) into C57BL/6NCrl mouse zygotes that were transferred to recipient female mice. Microinjection was performed at the Biocenter Oulu Transgenic and Tissue Phenotyping Core Facility, University of Oulu, Finland. Genomic DNA was isolated from earmark biopsies taken at weaning. Biopsy samples were lysed with 0.1 mg/ml proteinase K in lysis buffer (0.1 M Tris pH 8.5, 5 mM ethylenediaminetetraacetic acid, 0.2% sodium dodecyl sulfate, 0.2 M NaCl) and, thereafter, genomic DNA was precipitated with absolute ethanol, washed with 70 % ethanol, dried, and dissolved in TE buffer (Tris-HCl and 0,05M EDTA, pH 8,0). KI-positive pups were identified by restriction digestion of PCR product, and the edit was confirmed by Sanger sequencing. Primer sequences are presented in Supplementary Table I. Two founders carrying the p.R24G variant were mated with wild-type C57BL/6NCrl mice to establish two separate mutant lines (A and B), and the F1 pups were analysed using the same techniques as in the case of founder mice. All experiments, unless otherwise stated, were conducted with homozygous Cln8^R24G^ KI mice and their wild type (WT) littermates were used as a control. All data shown in the main manuscript was derived from the A line, and data presented in the Supplementary file was derived from A and B lines.

Animal care was carried out with the support of the Oulu Laboratory Animal Centre Research Infrastructure, University of Oulu, Finland. Animal care and experimental procedures were conducted in accordance with the national legislation, EU Directive 2010/63/EU, and the ethical standards of the institution. All animal procedures were approved by the Project Authorisation Board at the State Provincial Office of Southern Finland (ESAVI/33743/2019 and ESAVI-17463-2022).

### RNA extraction and cDNA synthesis

RNA extraction and cDNA synthesis were performed from liver and brain samples of Cln8^R24G^ KI mice and littermate controls. The RNA was extracted from the flash frozen left hippocampal part (three Cln8^R24G^ KI and three Cln8^R24G^ WT mice at 8 weeks of age, mixed sexes) using the RNeasy® Lipid Tissue Mini Kit (Qiagen, Venla, Netherlands) according to manufacturer’s instructions. The RNA extraction from flash frozen liver samples was performed using the Monarch total RNA miniprep kit (New England Biolabs, Ipswich, Massachusetts, USA), including the additional on-column DNase digest according to manufacturer’s instructions (four Cln8^R24G^ KI and four littermate controls at 8 weeks of age, mixed sexes). cDNA synthesis was performed using the QuantiTect Reverse Transcription Kit (Qiagen, Venla, Netherlands) according to manufacturer’s instructions. Samples were incubated for 30 minutes at +42 degrees after the reverse transcriptase master mix was added. To confirm the presence of the knock-in on RNA level, Sanger sequencing of the cDNA was performed as described above.

Primer sequences are presented in Supplementary Table I.

### Quantitative PCR

The *Cln8* RNA expression in the brain and liver was studied using qPCR. qPCR was performed according to the manufacturer’s instructions (IQTM SYBR Green Supermix, Bio-Rad, Hercules, CA, USA) using CFX ConnectTM Real-Time System (Bio-Rad, Hercules, CA, USA). Tm of the analysis was adjusted at 60°C. Ribosomal protein L13a (*Rpl13a*), hypoxanthine phosphoribosyltransferase 1 (*Hrt1*) and phosphoglycerate kinase 1 (*Pgk1*) were used as reference genes. The primers were designed using NCBI primer blast (Ye et al., 2012) (Supplementary Table I). To analyze the relative difference in the expression of *Cln8* between the KI and WT mice, the Paffl method was used. The standard deviation, standard error, and mean were calculated in GraphPad Prism and the statistical significance was determined using the Welch’s t-test in GraphPad Prism.

### Health monitoring

Systematic follow up of the homozygous Cln8^R24G^ KI mice and their littermates was started at 3 weeks of age and ended at 16 weeks (hereafter, 4 months) or at 36 weeks of age. Mice were euthanized at 37−38 weeks (hereafter 9 months) of age. Animals were kept on 12-h light/dark cycle and had unlimited access to chow (Teclad 2018/2918, Inotiv) and water. The bodyweight, physical appearance, and well-being of the animals was assessed once a week until 18 weeks of age and thereafter once every 2 weeks until the end of the experiment. If there was a significant change in the parameters being followed, the monitoring period was shortened or follow-up was ended prematurely, and the animal was euthanized. Health monitoring was carried out with the support of the Oulu Laboratory Animal Centre Research Infrastructure, University of Oulu, Finland. At the end of the experiment, mice were euthanized using carbon dioxide sedation and cervical dislocation. Tissues were collected for biochemical and histological analysis. Both sexes were used in the experiment.

### Gait analysis

CatWalk XT system (Noldus Information Technologies, the Netherlands) was used for gait analysis. Briefly, the mice were allowed to freely run in a corridor (length 1.5 m, width 5 cm) on a glass plate and illuminated paw prints were captured with a high-speed digital camera. First, a mouse with an average weight among all was used to calibrate the exposure settings (automatic). One mouse was heavier than others and required manual setting to optimize the footprints. Three acceptable runs were recorded from each mouse. The criteria were the following: The passage must be between 0.6–5.0 s and the maximum speed variation 50%. The glass surface of the corridor was wiped with window cleaning spray between each mouse. The system automatically annotated the footprints (left front and hind paw, right front and hind paw) correctly in most cases, but the researcher verified the classification and made manual corrections if needed.

For the analysis, the following parameters were selected: running speed and mean print area, mean swing duration, mean swing speed, base of support (lateral distance between the right and left paw) of the fore- and hind paws.

### Histopathological analysis

Tissue samples were collected 4 or 9 months of age. In the case of Cln8^R24G^ B line, the samples were collected at 33 or 36 weeks of age (see Supplement).

Brain and eye samples from mice, not included EEG recordings, were immersion fixed for 24h in either 4% paraformaldehyde in PBS (at +4℃, subset of brain and eye samples), 10% neutral buffered formalin [at room temperature (RT), subset of brain samples] or Davidson fixative (RT, subset of eye samples). The histology samples were washed for 1h under running tap water and stored in 70% ethanol until paraffine embedding. The tissue processing, using Tissue-Tek VIP® 6 AI Vacuum Infiltration Processor (Sakura, Torrance, CA, USA), was performed by the Transgenic and Tissue Phenotyping Core Facility at the University of Oulu. The paraffin-embedded brain samples were sectioned into 5µm sections corresponding to bregma 1.33 to 0.97 mm (striatum samples), bregma –1.43mm to –2.27mm (Hippocampus samples) and bregma –5.91mm to –6.23mm (Cerebellum samples) using the Epredia HM335s microtome (Portsmouth, New Hampshire, USA). Tissues from four mice of each genotype and sex for the 9-month timepoint, four individuals of each genotype for 4-month-old males, and three mice for each genotype for the 4-month-old females were used for the histological analysis of the brain. Unless otherwise indicated, paraffin-embedded eye blocks from mice aged 2, 4, and 9 months were sectioned at a thickness of 5 μm using a microtome (Thermo Scientific microm HM355S, Waltham, MA USA).

Mice that underwent electroencephalography (EEG) recordings and subsequent FosB staining were deeply anesthetized with pentobarbital/chloralhydrate cocktail (40 mg/kg each) and perfused through the heart, first with ice-cold saline and then with 4% paraformic acid solution. All perfusions were done in the afternoon between 1:00 – 3:00 p.m. The brain was extracted from the skull and further immersion fixed with 4% paraformaldehyde solution for 4 h followed by 30% sucrose overnight. The brain was stored in an antifreeze solution at −20°C until cut with a freezing slide microtome (Leica SM2000R) into 35 µm coronal sections.

The presence of autofluorescent storage material in the paraffin sections of the hippocampus and cerebellum was investigated as follows. 5µm sections were baked for 1h at 60°C. Subsequently, the samples were deparaffinized with two changes of 100% xylene (3−5 seconds each). The rehydration was done by dipping the slides for 5−10 seconds into two changes of ethanol absolute, one change of 95% ethanol and one change of 70% ethanol. The samples were then washed once in sterile water and then twice with 1x PBS, and counterstained with 1 µg/ml DAPI (Merck, Rahway, New Jersey, USA) for 20 minutes at RT. The sections were washed 3x with 1xPBS and mounted with Immu Mount (Epredia, Portsmouth, NH, USA). The samples were left to dry overnight and scanned using the Olympus Slideview Vs200 universal whole slide imaging scanner (Tokyo, Japan). The excitation wavelengths for the imaging were 405nm for DAPI and 488nm for autofluorescence.

For the Luxol Fast Blue staining, sections were baked, deparaffinized by two changes of 100% xylene and then rehydrated in a descending alcohol line until 95% ethanol. The slides were then transferred to prewarmed (60°C) and filtered Solvent Blue 38 staining solution [0.1% Solvent Blue 38 (Sigma Aldrich St. Louis, Missouri, USA) and 0.05% acetic acid (Fisher chemicals, Pittsburgh, Pennsylvania, USA) in 94% ethanol] and incubated for 3 hours at 60°C. The slides were subsequently washed in distilled water for 30−45 seconds and then differentiated in 0.5% aqueous lithium carbonate solution (Sigma Aldrich St. Louis, Missouri, USA) for 5 seconds. After differentiation, the samples were immediately washed in distilled water. For the counterstaining, the samples were incubated in Harris hematoxylin (Leica, Nussloch, Germany) and blued under running tap water. Samples were dipped two times into 94% ethanol and then transferred to eosin (Sigma Aldrich Sigma Aldrich St. Louis, Missouri, USA). The samples were washed in two changes of 70% ethanol and then incubated in two changes of ethanol absolute. Finally, the samples were briefly transferred to 100% xylene and cover slipped with Tissue-Tek® coverslipping film using the Tissue-Tek Film® automated coverslipper (Sakura, Torrance, CA, USA). Selected tissue slides were scanned at the Biocenter Oulu Transgenic and Tissue Phenotyping core facility with NanoZoomer S60 digital slide scanner (Hamamatsu, Hamamatsu City, Shizuoka Prefecture, Japan).

### Immunohistochemistry

Immunohistochemical (IHC) stainings in brain samples (with the exception of FosB stainings) were performed using the ImmPRESS HRP Horse anti-rabbit IgG plus polymer kit (MP-7801, Vector labs, Newark, CA, USA) and different primary antibodies (GFAP (E4L7M) XP#80788, Cell Signaling Technology, dilution 1:400; CD68 (E3O7V) Rabbit mAb #97778, Cell Signaling Technology, dilution 1:300; NeuN (D4G4O) XP® Rabbit mAb #24307, Cell Signalling Technology, dilution 1:800 and Anti-ATP synthase C antibody [EPR13907] ab181243, Abcam, dilution 1:2000) were used to visualize the accumulation of storage material, neuroinflammatory processes as well as neuronal loss. The staining protocols were adapted from the manufactures’ instructions. The demasking was done by boiling the samples in 0.01M citrate buffer; pH 6.3%, hydrogen peroxide was used for quenching the endogenous peroxidases for 10 minutes at RT. The primary antibodies were diluted in 1x PBS and the samples were incubated with the antibody dilutions overnight at +4℃. For washing steps, 1X PBS containing 0,1% (v/v) Tween 20 was used. The samples were counterstained using Harris hematoxylin (Leica, Nussloch, Germany), blued in running tap water, and thereafter dehydrated, cover slipped, and scanned as described above.

FosB immunohistochemical staining of brain sections from mice that underwent EEG recordings was prepared by pre-treating them with 0.05 M citrate solution (pH 6.0) in an 80°C water bath for 30 minutes. The brain sections were allowed to cool down in 0.5 M Triton/Tris buffered saline (TBS-T, pH 7.6), after which the sections were washed 3×5 min in 0.5 M TBST, pH 7.6. To analyse long-term neural activity, the sections were stained with anti-rabbit-FosB (1:5000, Abcam; ab184938) at +4 °C overnight. The sections were incubated in goat-anti-rabbit AlexaFluor 594 (1:400, Invitrogen, A-11012) for two hours at RT. The sections were mounted with Mowiol® 4-88 and left to dry RT until imaging. Brain sections stained with FosB immunohistochemistry were imaged with Zeiss Image r.M2 fluorescence microscopy at the University of Eastern Finland using Zen 3.3 software.

### Transmission electron microscopy

Transmission electron microscopy (TEM) was performed at the Biocenter Oulu Electron Microscopy Core Facility. For TEM analysis, brain slices were fixed in 1% glutaraldehyde, 4% formaldehyde mixture in 0.1 M phosphate buffer for 12−16 hours, post-fixed in 1% osmium tetroxide, dehydrated in acetone, and embedded in Epon LX 112 (Ladd Research Industries, Williston, Vermont, USA). Thin sections (70 nm) were cut with an ultramicrotome (FC6; Leica, Wetzlar, Germany), stained in uranyl acetate and lead citrate, and examined in Tecnai G2 Spirit 120 kV transmission electron microscope (FEI Europe, Eindhoven, The Netherlands). Images were captured by Quemesa CCD camera and analysed using Radius software (EMSIS GmbH, Munster Germany).

### Quantification of staining

Quantification of NeuN -positive and -negative cells in the brain of Cln8^R24G^ mice was performed using Visiopharm software (Visiopharm, Hoersholm, Denmark). The regions of interest (ROIs) included the thalamic nuclei (in particular VPN/VPL) and the somatosensory cortex barrel field 1 (SB1F) in 4-month and 9-month-old Cln8^R24G^ KI mice and littermate controls (both sexes). Each group included 3 individuals, with exception of 4-month-old females, which comprised only two Cln8^R24G^ KI mice and two littermate controls. ROIs were manually drawn. The AI was trained using images from two 4-month-old mice for each genotype and sex. Training data was generated utilizing the analysis protocol baggage (APP) 10167 - Nuclei Detection, AI (Brightfield) for nuclei annotation, followed by manual correction and outlining to aid segmentation. The above-mentioned APP was customized for outputs and through training (and assessed on brain sections of 4-month-old Cln8^R24G^ mice for accuracy). The left and right hemispheres were quantified separately. The analysis included the quantification of NeuN-positive and NeuN-negative nuclei for each region of interest (ROI). Nuclear number was counted and normalized to ROI area to get the nuclear density value, which was compared between groups. The statistical analysis was performed in groups of mice containing more than two individuals using GraphPad Prism (including standard deviation, standard error, and mean). Welch’s t-test was performed to assess statistical significance in groups containing more than two individuals and was omitted in groups containing two or less individuals in this experiment.

FosB intensity was quantified in the dentate gyrus in two sections, at −1.7 to −1.9 and at −2.5 to −3.1 from bregma using ImageJ software. First, the mean grey value was measured from the granule cells layer bilaterally and averaged. Second, the background mean grey value was measured from the cell free molecular layer of the lower blade of two dentate gyri, averaged, and subtracted from the signal. The FosB intensity between the genotypes was compared using nonparametric Mann-Whitney U-test.

### EEG analysis of seizure activity

#### Electrode implantation

The mice were implanted with triple wire electrodes in the left hippocampus, and with conventional skull screw electrodes bilaterally above the frontal cortex. Screw electrodes above the cerebellum worked as the ground and reference as well as anchors for dental acrylic cement for a miniature connector. Electromyogram (EMG) was recorded with a stainless-steel wire between the neck muscles. Under isoflurane anaesthesia (4.5%–2%), two cortical screw electrodes were attached to the left and right frontal bone (AP +1.7, ML +/– 1.8 from bregma). A bundle of three 50 µm insulated stainless-steel wire electrodes (stagger 350 µm between the tips) was installed into the left and right hippocampus (AP −2.2, ML +1.5 from bregma, DV 1.3 from dura). The location of the deep electrodes was confirmed using electrophysiological landmarks as well as histologically. Carprofen (5 mg/kg, i.p.) was given for postoperative analgesia. The mice were given one week of recovery time after the operation before they were familiarized with the recording environment.

#### Video EEG acquisition and data analysis

The video EEG recording took place in a circular frame made of brown hardboard (diameter 18.5 cm, wall height 18 cm) on a translucent glass plate that was illuminated from below. Each mouse was recorded individually. A light recording cable with a counterweight through a pulley system was attached, fixed to a preamplifier (Plexon) that connected to the imbedded connector on the mouse head. The other end of the recording cable was connected to an AC amplifier (AM Systems; gain 1000, analogue band-pass 1-3000 Hz). The amplified signal was digitized at 2 kHz per channel. The movements of the animals were recorded with an overhead video camera. Synchronized EEG and video signals were acquired using the SciWorks 5.0 program (A-M Systems). Each recording session took 3 h, either in the morning or afternoon, but always during the light period. The lights were on in the laboratory throughout the recording. The mice were adapted to the laboratory environment and plugging of the recording cable for two weeks before the recording onset and spent most of the recording session in sleep.

Separation between movement and immobility was based on the EMG. We band-pass filtered EMG at 1−100 Hz and calculated its envelope. The envelopes clearly show periods of low and high amplitude, the former representing moving epochs while the latter correspond to immobility. Further, if the mouse moved <0.5 s between two brief immobility epochs or <1 s between two immobility epochs of >5 s, this short movement was assigned to immobility. When immobility lasted >20 s, the first 20 s were defined as waking immobility, the rest as sleep. Furthermore, we searched for REM in sleep epochs with low delta (1−4 Hz) and high theta (6−9 Hz) amplitudes in the top hippocampal channel. First, we searched for epochs where delta or theta envelope was >100 mV for >400 ms and connected two epochs if the gap in between was <1 s. Epochs with only theta envelope above these thresholds was defined as REM. If the gap between two REM epochs was <10 s, they were connected. The rest of sleep time was defined as NREM. Finally, the total sleep duration was calculated as the sum of NREM and REM epochs.

The spike extraction from the EEG was done with a customized Matlab program. First, after 50 Hz notch filtering, all channels were high-pass filtered at 8 Hz to remove slow fluctuations from the baseline. First, the hippocampal channel with the highest deviation from baseline was selected and peaks >8 SD from the baseline and <50 ms in duration were extracted. The hippocampal spikes were considered epileptiform discharges (ED) if a sharp peak was seen in all channels (one cortical channel >3 SD) and was followed by after-hyperpolarization lasting >200 ms. Since epileptiform spiking in APP/PS1 mice mostly happens during sleep, the detected ED counts were normalized to the total sleep time and expressed and EDs/h.

### Retinal IHC and microscopy

The IHC for eye samples from Cln8^R24G^ KI mice and littermate controls was conducted according to the following protocol. The sections were deparaffinized in xylene (3×5 min) and ethanol series (99.9% 2×5 min, 94% 1×5 min, 70% 1×5 min, and 50% 1×5 min), rehydrated in MQ water (1×1 min), and washed in tris-buffered saline with or without Triton to increase tissue permeability (TBS, pH 7.5, 1×5 min; TBST 0.1%, 1×5 min). The sections were blocked with 2% BSA (Bovine Serum Albumin) prepared in TBST 0.1% for 1 hour and then incubated in primary antibodies (GFAP (GA5) mouse mAb #3670, Cell Signalling Technologies, dilution 1:2000; Rabbit anti-opsin1 Rb0216-25-1107-w8, NOVUS, dilution 1:1000), with mild shaking overnight at RT. After that, the sections were washed for 3×5 mins in TBST 0.1% and incubated with a fluorescent secondary antibody (AF donkey anti-mouse 568nm, ab175472, Abcam, dilution factor 1:500; and AF donkey anti-rabbit 647nm ab150075, Abcam, dilution factor 1:500) for 2 hours in the dark, at RT with mild shaking. Each slide contained one negative control sample. Finally, the slides were washed again for 3×5 mins in TBST 0.1% and 1×5 mins in TBS to ensure removal of residual detergent from the glass surface. The sections were thoroughly air-dried for approximately 30 mins and subsequently mounted using fluoroshield mounting medium with DAPI (ab104139, Abcam).

Fluorescent imaging of the sections was performed using an Olympus APEXVIEW APX100 microscope equipped with an Olympus U-LGPS light source at 20× magnification. All images were acquired from the central region of the retina.

## Results

### Cln8^R24G^ KI mice develop progressive motor symptoms and spontaneous seizures

To study disease mechanisms related to Northern epilepsy CRISPR-Cas9 gene editing was used to generate two Cln8^R24G^ founder lines (A and B). Due to the lack of a specific antibody against Cln8, validation was restricted to the genomic and RNA levels. Sanger sequencing confirmed the presence of the p.R24G variant and the silent variant that produced an *Nhe*I restriction site in the line A mutant mice but not in the littermate controls (Figure 1A and B). In addition to the p.R24G variant, mouse line B carried two silent variants, one creating the *Nhe*I restriction site, as in line A, and another designed to prevent re-cutting by S.p. HiFi Cas9 in the target region (Supplementary Figure 1).

**Figure 1.**
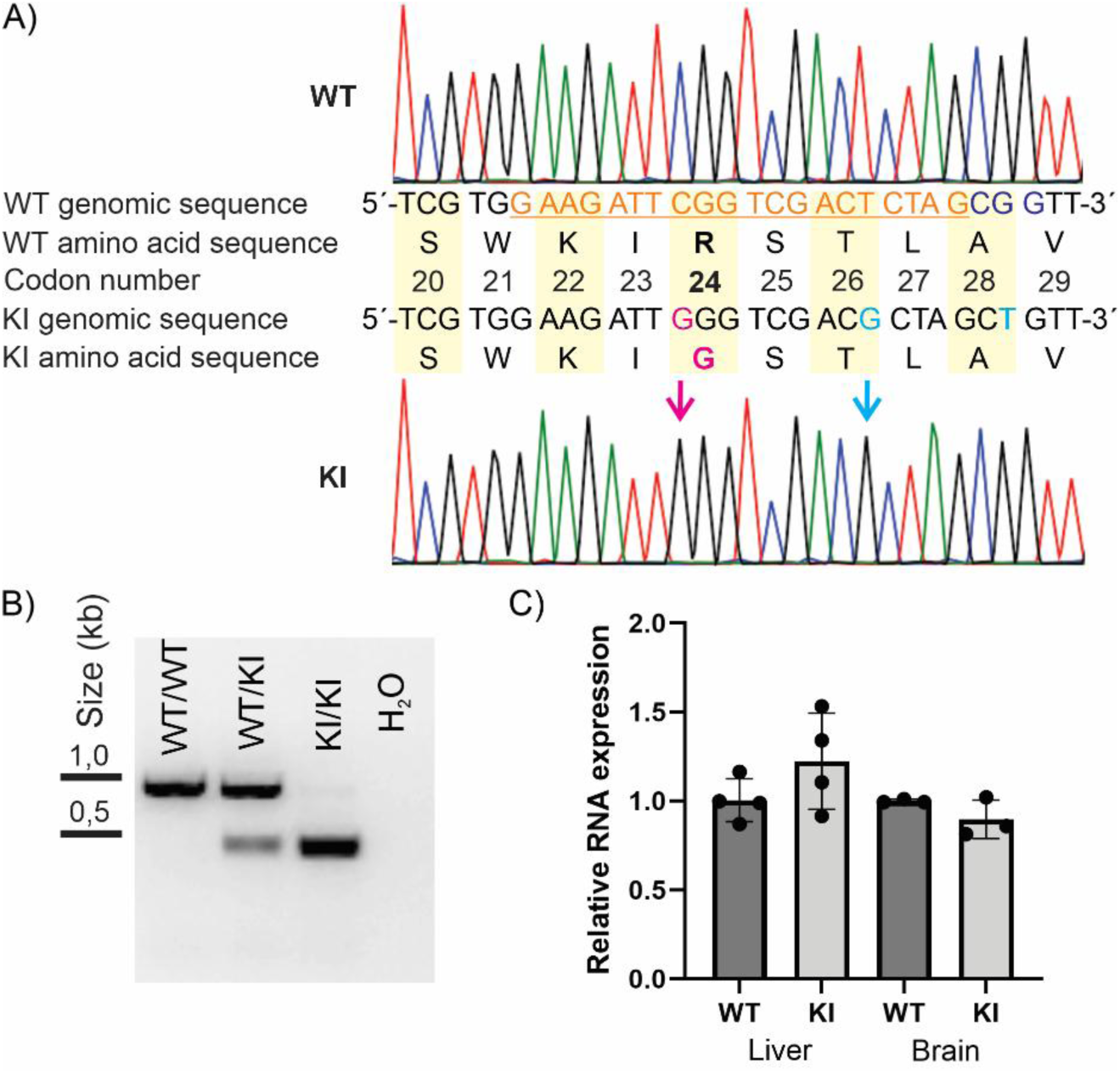
Validation of the Cln8^R24G^ KI at the genomic and RNA levels. A) Sanger sequencing results showing the p.R24G variant and a silent variant introducing an *Nhe*I restriction site. In the WT genomic sequence, the guide sequence is underlined and highlighted in orange, and the PAM sequence is highlighted in blue. B) Agarose gel electrophoresis of DNA fragments after *Nhe*I digestion showing different genotypes. *Nhe*I digestion of the KI allele yielded 470-bp and 421-bp products, whereas the WT allele remained undigested (891 bp). C) Relative *Cln8* RNA expression in the liver (*p* = 0.21) and the brain (*p* = 0.18927) of the WT and KI mice. The qPCR analysis was conducted using tissues from 2-month-old mixed-sex mice (brain: WT *n* = 3, KI *n* = 3; liver: WT *n* = 4, KI *n* = 4). Abbreviations: knock-in, KI; wild type, WT; kilobase, kb.

Quantitative RT-PCR was used to analyse the effect of the p.R24G variant on RNA expression. No significant differences were detected between *Cln8* mRNA expression in the liver and brain of the Cln8^R24G^ KI mice and that of the littermate controls indicating that the amino acid substitution did not alter the *Cln8* mRNA levels (Figure 1C).

The Cln8^R24G^ KI mice were viable at birth and asymptomatic. To characterise the phenotypic spectrum associated with the Northern epilepsy variant, systematic health monitoring was conducted on 6 WT and 5 KI males and 8 WT and 7 KI females assessing body weight and general well-being, from 3 weeks of age (weaning) up to 9 months. The mice were group-housed under a 12-h light/dark cycle with ad libitum access to chow and water. At approximately 7 months, the Cln8^R24G^ KI mice began to develop progressive motor symptoms. Three female Cln8^R24G^ KI mice suddenly died and 1 male and 1 female were euthanised due to deteriorating health before the end of the monitoring period. The euthanised female Cln8^R24G^ KI mouse displayed a clear seizure with noticeable muscle jerks. The male Cln8^R24G^ KI mice showed weight loss from 7 months onwards, although this was not statistically significant, while the female Cln8^R24G^ KI mice did not show weight loss (Figure 2A). Six of the seven Cln8^R24G^ KI mice that survived to the end of the monitoring period exhibited motor symptoms ranging from mild hind-leg clumsiness to hind-leg paralysis. The KI mice also developed eye discharge and periocular hair loss (Figure 2B). At 9 months, in one KI mouse twitches/jerks resembling myoclonic seizures −a key feature of Northern epilepsy− were observed. Similar symptoms were subsequently noted in several Cln8^R24G^ KI mice from mutant lines A and B.

**Figure 2.**
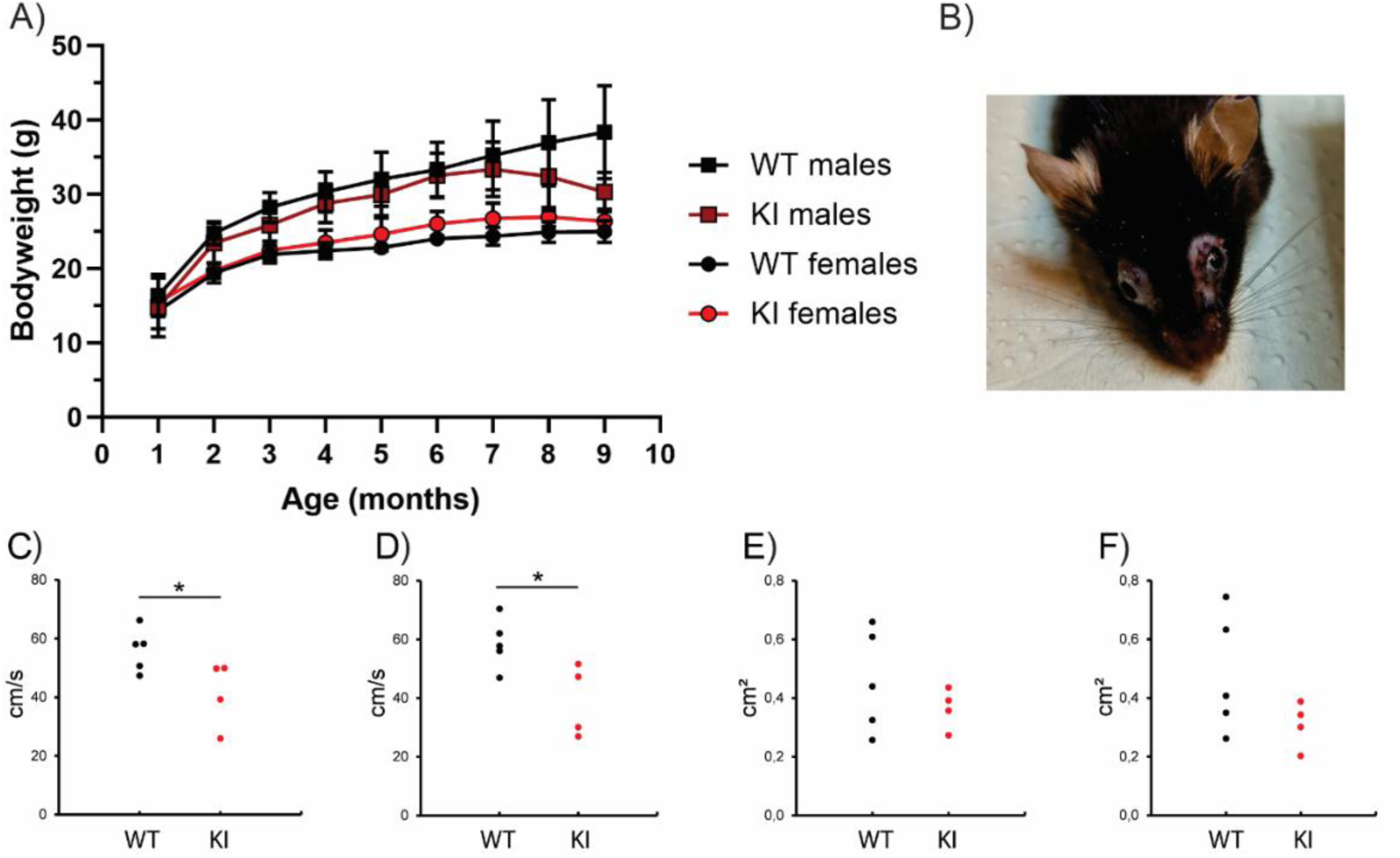
Health monitoring and catwalk gait analysis of Cln8^R24G^ mouse line. A) Bodyweight of the male and female Cln8^R24G^ KI mice and their littermates from 3 weeks to 9 months of age (males: WT, *n* = 6; KI, *n* = 5 of which 4 survived to the end of the experiment; females: WT, *n* = 8; KI, *n* = 7 of which 3 survived to the end of the experiment). Data points are shown for full months. B) Female Cln8^R24G^ KI mouse showing hair loss and eye discharge. Similar problems were observed in 6 of the 7 Cln8^R24G^ KI mice at the end of the follow-up period. Catwalk gait analysis: C) Mean swing speed of the front paws, D) mean swing speed of the hind paws, E) mean print area of the front paws, F) mean print area of the hind paws. \**p* < 0.05, Student’s t-test. Abbreviations: knock-in, KI; wild type, WT.

To evaluate motor function, gait analyses were performed on 5 WT and 4 Cln8^R24G^ KI mice at 7 months. The main findings from the gait analysis are shown in Figure 2C-F. No genotype differences were observed in the static parameters, including the paw print area and the base of support for the front and hind paws (Figure 2E and F). The running speed was marginally slower in the Cln8^R24G^ KI mice (*p* = 0.06), and the swing speed of both the front and hind paws was significantly slower (p < 0.05) (Figure 2C and D).

### Cln8^R24G^ brains accumulate autofluorescent, SCMAS and LFB -positive storage material with curvilinear-like ultrastructures

Histological analyses of the central nervous system were performed with emphasis on the striatum, hippocampus, thalamic nuclei, cortex and cerebellum, focusing on putative autofluorescent accumulations and their composition and distribution.

No gross anatomical abnormalities were observed in the Cln8^R24G^ KI mouse brains. Granular autofluorescent storage material was present in the hippocampus, thalamic nuclei, cortex, Purkinje cells and cerebellar nucleus at 4 months (Figure 3A and B, Figure 4A and B respectively). Accumulation progressed and was markedly increased by 9 months in all the Cln8^R24G^ KI brain regions analysed compared with the littermate controls (Figure 3C and D, Figure 4C and D).

**Figure 3.**
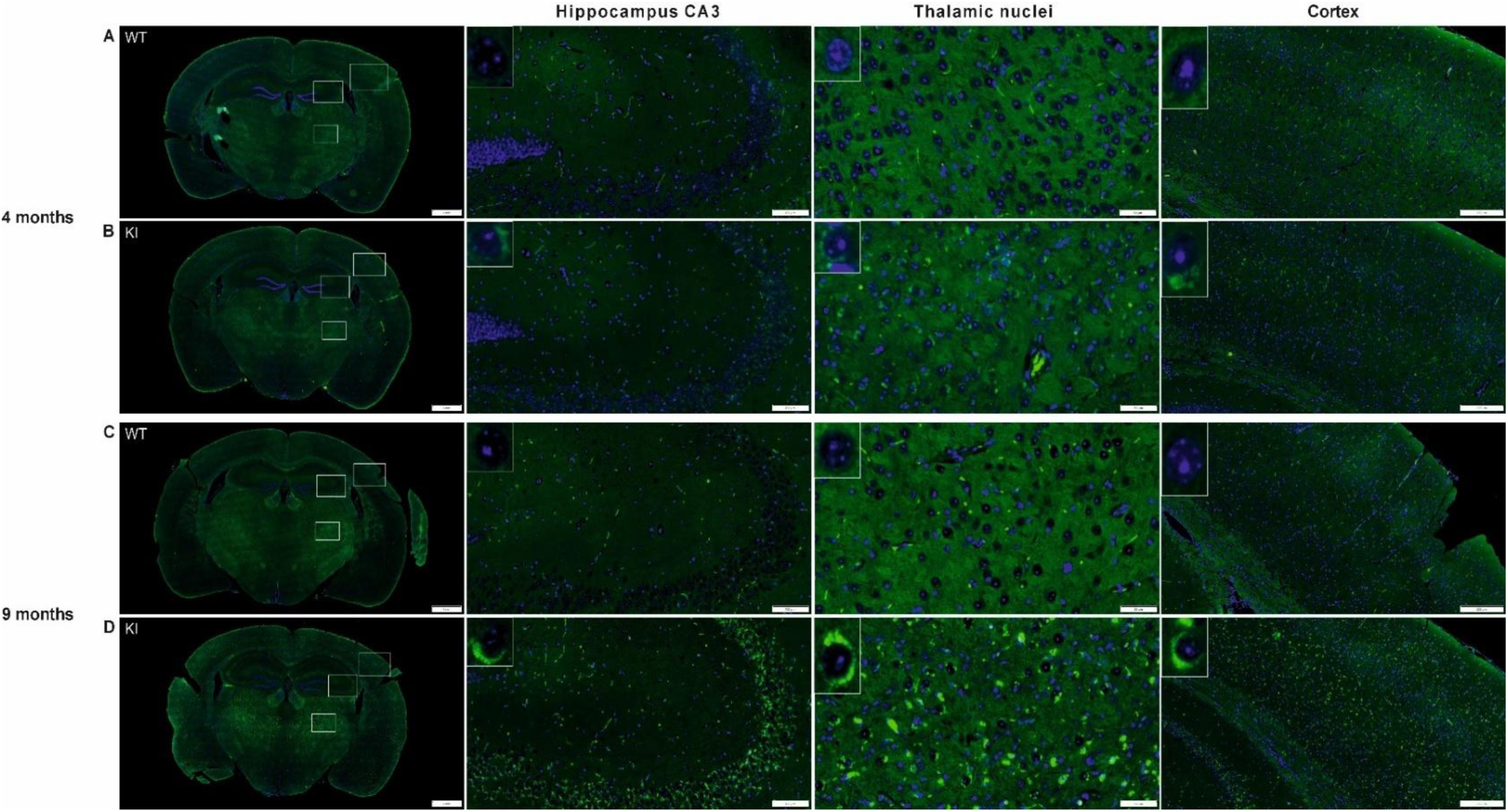
Accumulation of autofluorescent material in the hippocampus, thalamic nuclei and cortex of Cln8^R24G^ KI mice. Accumulation of autofluorescent material (at 488 nm excitation, green signal) was evident as early as 4 months of age in the Cln8^R24G^ KI mice (B). The accumulation increased with age (D), while no changes were detected in the littermate controls (A and C). The accumulation was most prominent in the hippocampus (especially CA3) and thalamic nuclei at 4 months, and markedly increased by 9 months. The autofluorescent material was also observed, though less extensively, in the cortex of Cln8^R24G^ KI mice. Representative images are from male mice (4 months: WT *n* = 3, KI *n* = 3; 9 months: WT *n* = 3, KI *n* = 3). Scale bars: overview, 1 mm; hippocampus, 100 µm; thalamic nuclei, 50µm; cortex, 200 µm.

**Figure 4.**
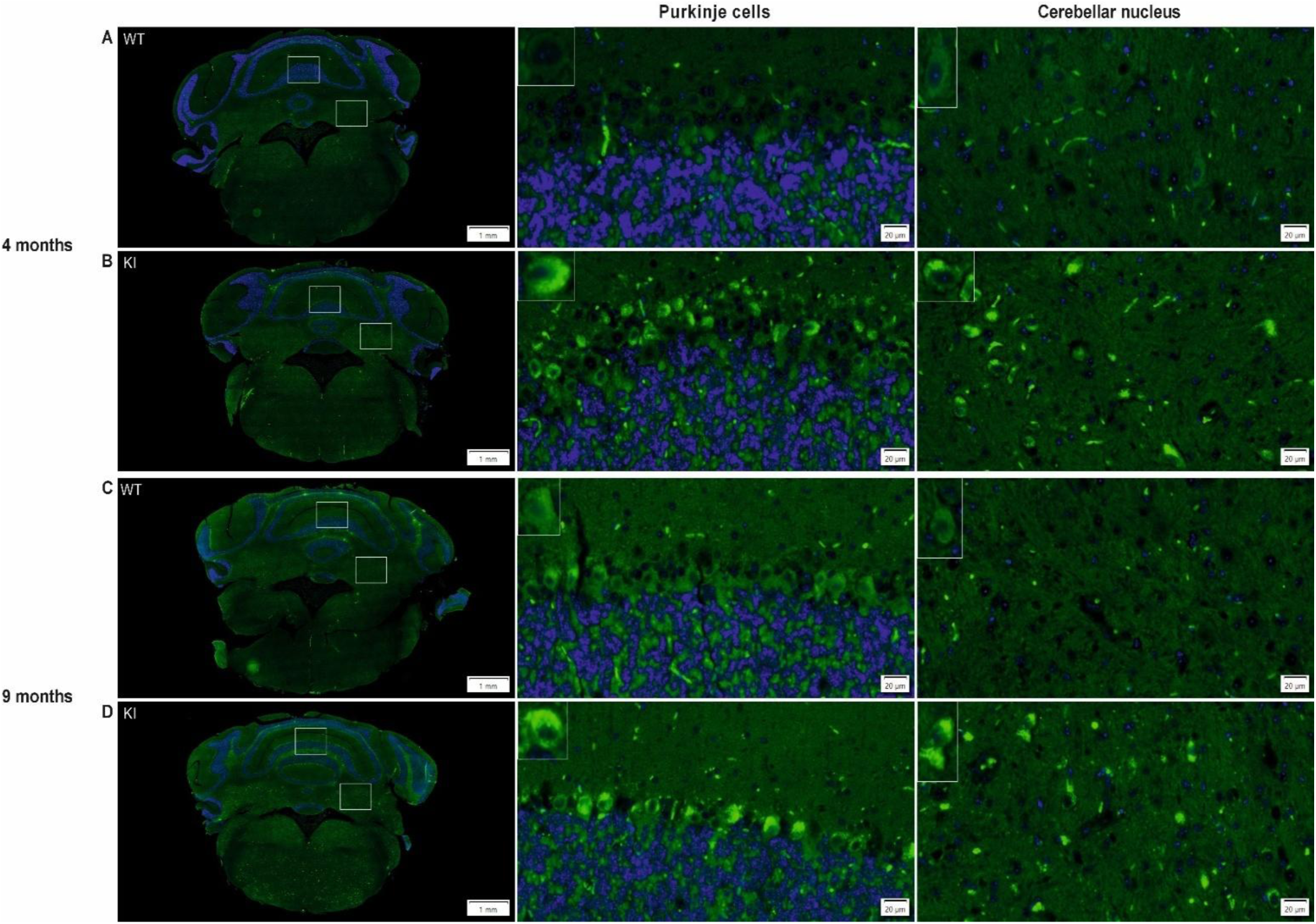
Accumulation of autofluorescent material in the Cln8^R24G^ KI cerebellum. Autofluorescence was evident in the Purkinje cells and cerebellar nucleus as early as 4 months of age (B) and progressively increased with age (green signal) (D). No autofluorescent accumulation was detected in control mice (A and C). Representative images are selected from male mice (4 months: WT *n* = 2, KI *n* = 2; 9 months: WT *n* = 3, KI *n* = 3). Scale bars: overview, 1 mm; Purkinje cells and cerebellar nucleus, 20 µm.

The distribution and staining characteristics of the autofluorescent material were further investigated using immunohistochemistry. SCMAS positive granular accumulations were evident in all the brain regions of the Cln8^R24G^ KI mice at 4 months (Figure 5B, 6B and Supplementary Figure 2B) and increased in intensity and number by 9 months (Figure 5D, 6D and Supplementary Figure 2D). The anterior commissure was largely SCMAS -negative, with occasional positive cells detected by 9 months (Supplementary Figure 2B, D). Accumulation of SCMAS was especially prominent in the hippocampal pyramidal cells, especially in the CA3 region (Figure 5B and D). Prominent accumulation was also observed in thalamic nuclei (especially VPL; Figure 5B and D), layer 5 pyramidal cells in the motor cortex (Supplementary Figure 2B and D) and the cerebellar nuclei (Figure 6B and D). At 9 months, Purkinje cells in the cerebellum showed marked SCMAS accumulation (Figure 6D), which was absent in the littermate controls (Figure 6C). Analysis of the hippocampus in 9-month-old females and male mice from founder line B line revealed a similar pattern (Supplementary Figures 3 and 4).

**Figure 5.**
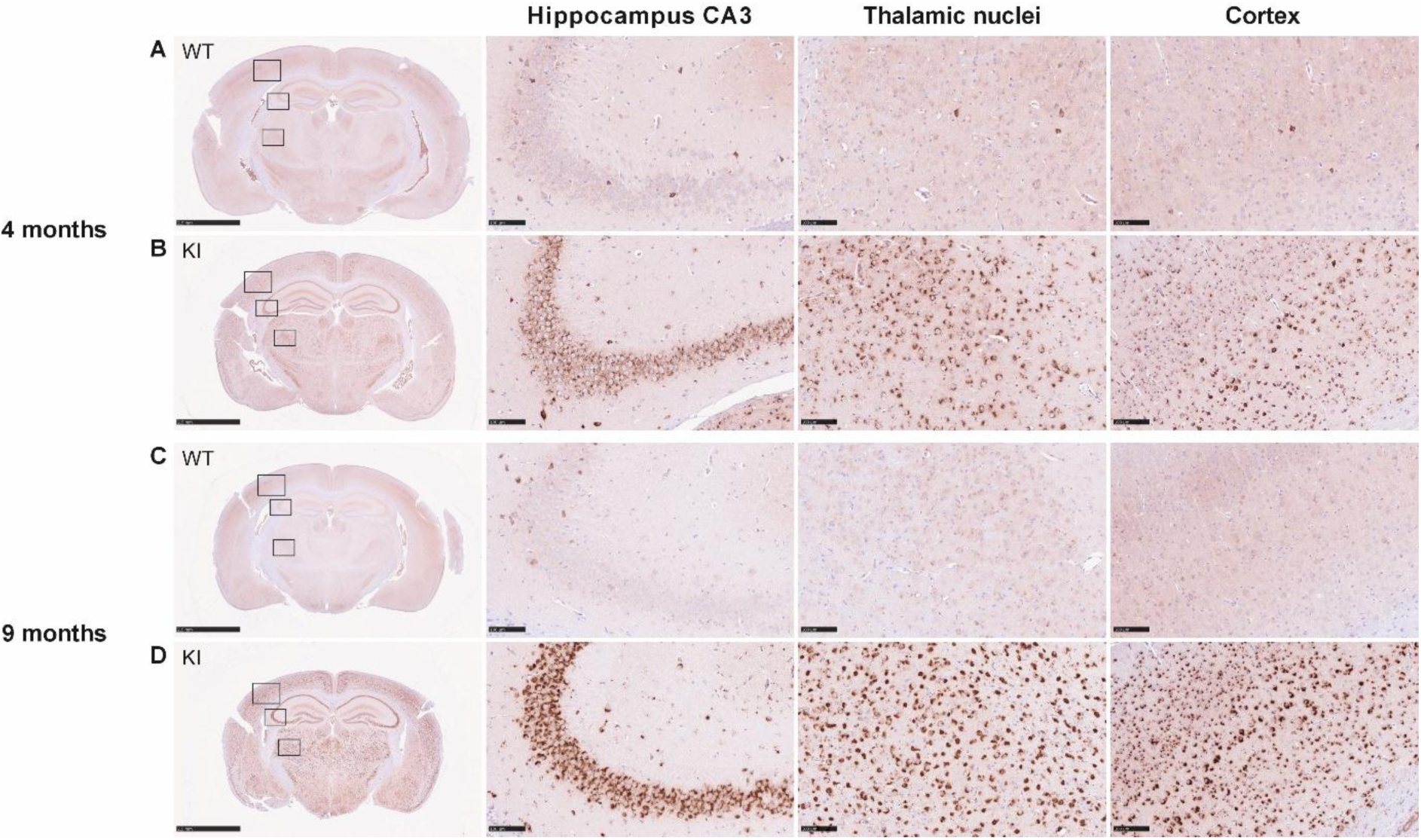
SCMAS staining of the hippocampus. Prominent granular staining, representing the autofluorescent accumulation, was present in the CA3 region of the hippocampus, thalamic VPL nucleus and somatosensory cortex of the Cln8^R24G^ KI mice at 4 months (B). Accumulation showed progressive increase with age (D). Minimal staining was observed in the littermate controls at the corresponding ages (A and C). Representative images are selected from male mice (4 months: WT *n* = 4, KI *n* = 4; 9 months: WT *n* = 4, KI *n* = 4). Scale bars: overview, 2.5 mm; hippocampus, thalamic nuclei and cortex, 100 µm.

**Figure 6.**
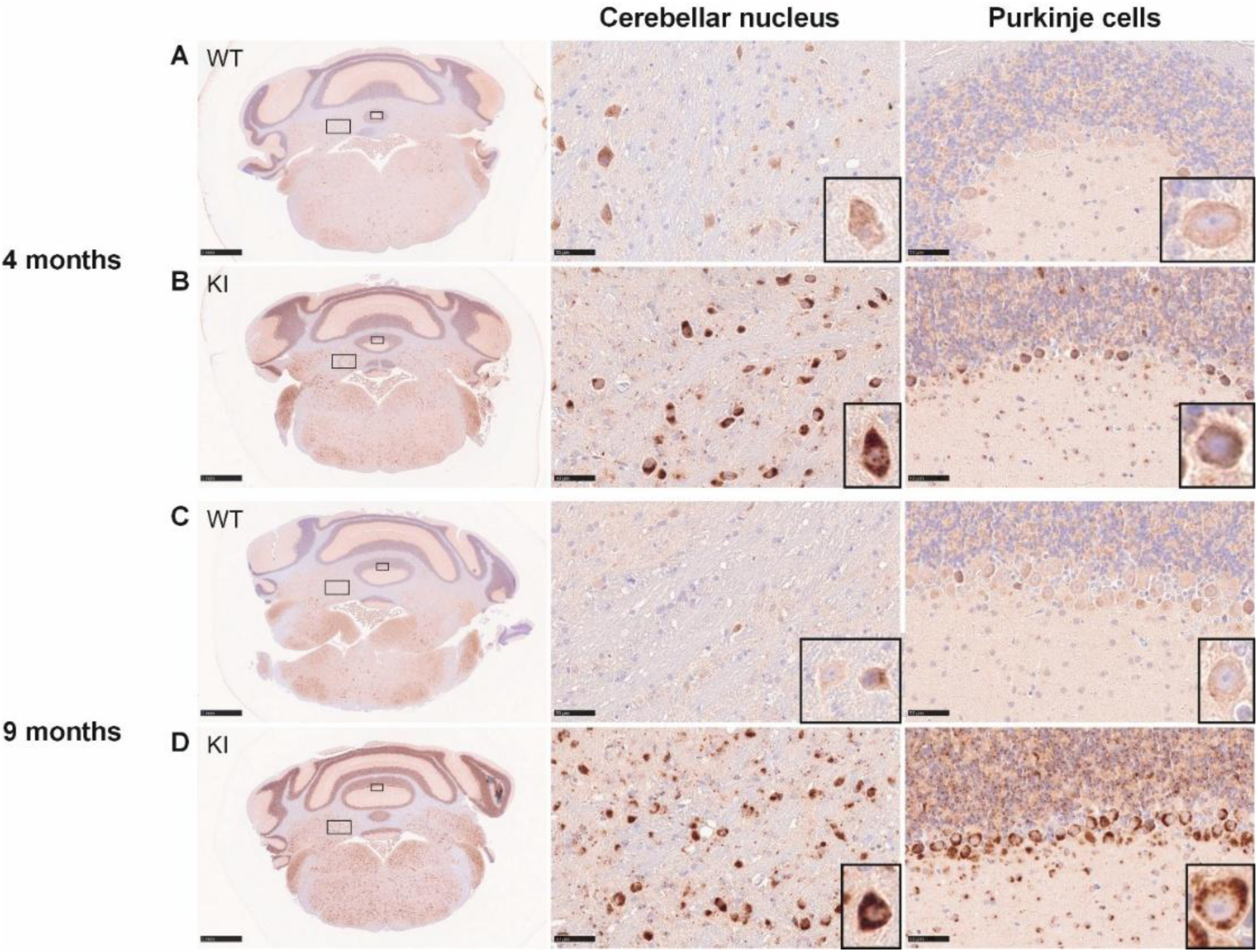
SCMAS staining of the cerebellum. Prominent granular SCMAS immunoreactive accumulation was evident in the Purkinje cells (zoomed images) of the Cln8^R24G^ KI mice (B and D). Occasional staining was also observed in the Purkinje cells of the littermate controls at both ages (A and C). The cerebellar nuclei of the Cln8^R24G^ KI mice (B and D), but not that of the littermate controls (A and C), showed marked accumulation. Representative images are selected from male mice (4 months: WT *n* = 2, KI *n* = 2; 9 months: WT *n* = 3, KI *n* = 3), but female mice were also analysed (data not shown). Scale bars: overview, 1 mm; cerebellar nucleus and Purkinje cells, 50 µm.

Ceroid lipofuscin, a characteristic feature of NCL neuropathology, has been shown to stain positively with Luxol Fast Blue (LFB) (Herva et al., 2000). In the Cln8^R24G^ KI mouse brains, several regions were strongly stained with LFB, with progressive accumulation of LFB-positive granules in the cortex, hippocampus and thalamic nuclei (Figure 7B and D). These structures were absent in the littermate control brains (Figure 7A and C). LFB also binds to phospholipid base-groups, including sphingomyelin, and is commonly used to stain myelin; however, no differences in myelination were observed between the littermate control and mutant brains at ages 4 and 9 months.

**Figure 7.**
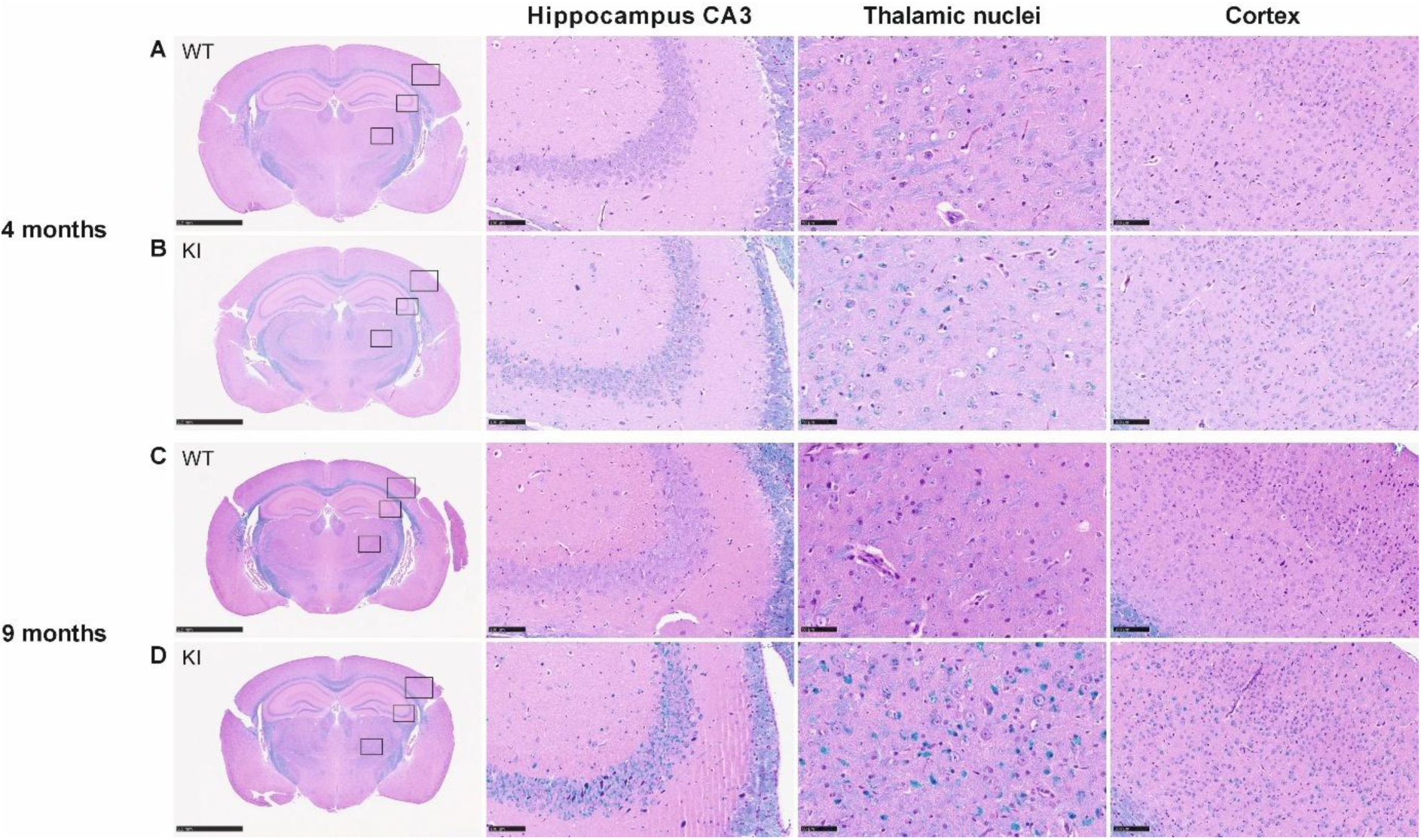
Luxol Fast Blue (LFB) staining in the hippocampus of Cln8^R24G^ KI mice and littermate controls at 4 and 9 months of age. An increase in LFB positive granules was evident in the hippocampus, thalamic nuclei and cortex of the Cln8^R24G^ KI mice as early as 4 months (B). The staining intensity increased with age, reflecting progressive accumulation (D). Similar LFB positive accumulation was not seen in littermate controls (A and C). Representative images are selected from male mice (4 months: WT *n* = 2, KI *n* = 2; 9 months: WT *n* = 3, KI *n* = 3). Scale bars: overview, 2.5 mm; hippocampus and cortex, 100 µm; thalamic nuclei, 100 µm.

The ultrastructure of ceroid lipofuscin in the brains of Northern epilepsy patients has been described as granular and curvilinear (Mole et al., 2005). Ultrastructural analysis of Cln8^R24G^ KI brains revealed electron-dense material with a curvilinear-like profile within cortical neurons at 4 and 9 months (Figure 8, panels on the right). Such deposits were absent in the wild type littermate samples (Figure 8, panels on the left). Lipofuscin deposits were occasionally seen in 9-month-old Cln8^R24G^ WT mice, but their ultrastructure differed markedly from those in the Cln8^R24G^ KI brains (Figure 8, bottom panels).

**Figure 8.**
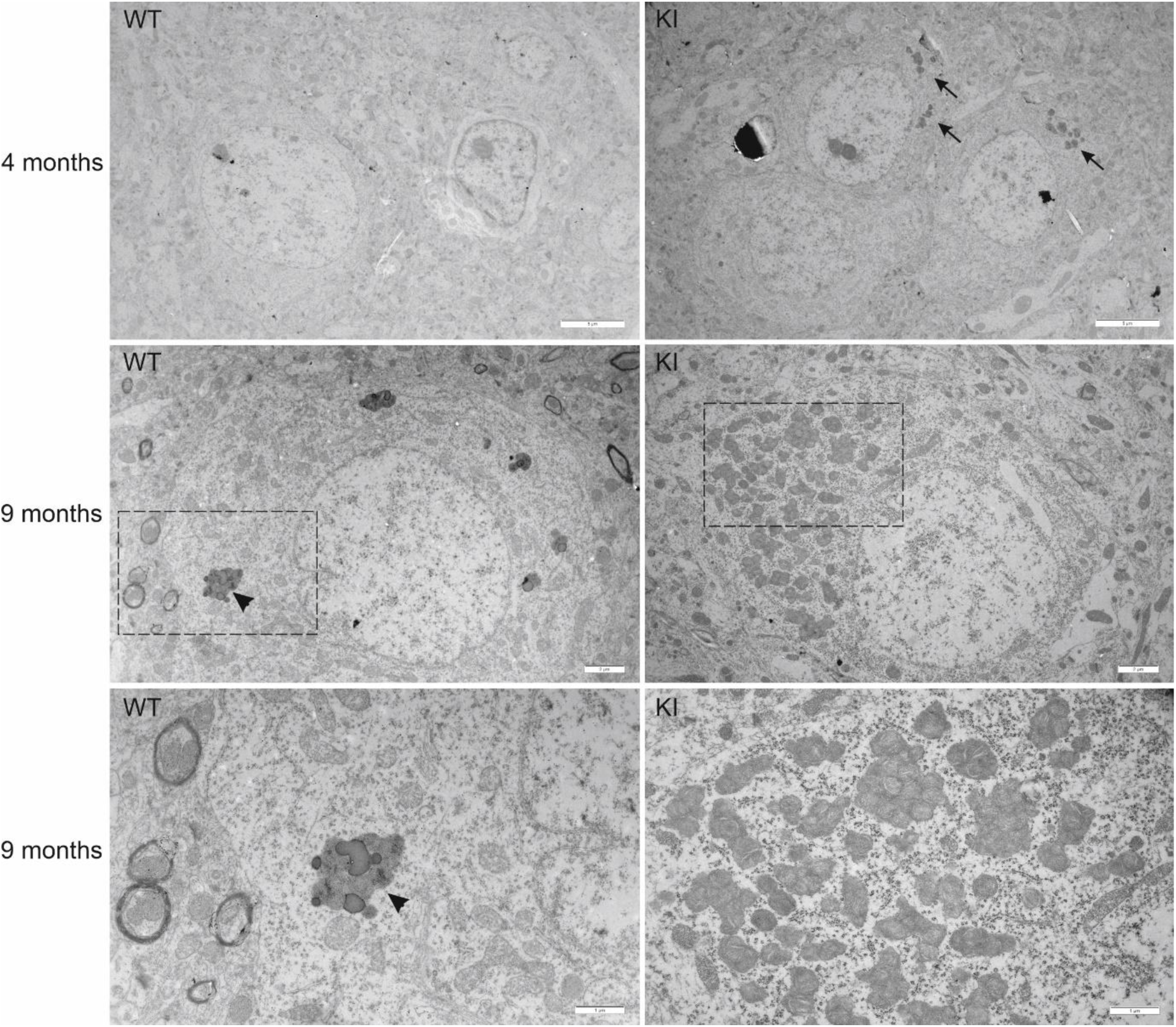
Ultrastructural analysis of accumulation in the cerebral cortex. Electron microscopy analysis revealed accumulation of electron-dense, curvilinear-like material within neurons of the cerebral cortex of 4- and 9-month-old Cln8^R24G^ KI male mice (panels on the right). Similar accumulation was absent in the cerebral cortex of the wild type littermates (panels on the left). Higher-magnification images (bottom panels) show lipofuscin deposits in cortical neurons of the WT mice (bottom left) and curvilinear-like ceroid lipofuscin deposits in cortical neurons of the Cln8^R24G^ KI mice (bottom right). Arrows indicate ceroid lipofuscin, and arrowheads lipofuscin. *n* = 3 in both genotype groups. Scale bars: 4 months, 5 µm; 9 months middle panels, 2 µm; and 9 months lower panels, 1 µm.

### Progressive neuroinflammation and significant neuronal loss in the thalamic VP nuclei of Cln8^R24G^ KI mice

Neuroinflammation was assessed using immunostaining for markers of activated astrocytes (GFAP) and microglia (CD68). Activation of both cell types was evident by 4 months (Figure 9B and Figure 10B) and increased markedly thereafter (Figure 9D and Figure 10D). GFAP positive (GFAP^+^) and CD68 positive (CD68^+^) cells were present throughout the sections and were especially prominent in the thalamic nuclei (VPN/VPL) (Figure 9B and D, Figure 10B and D).

**Figure 9.**
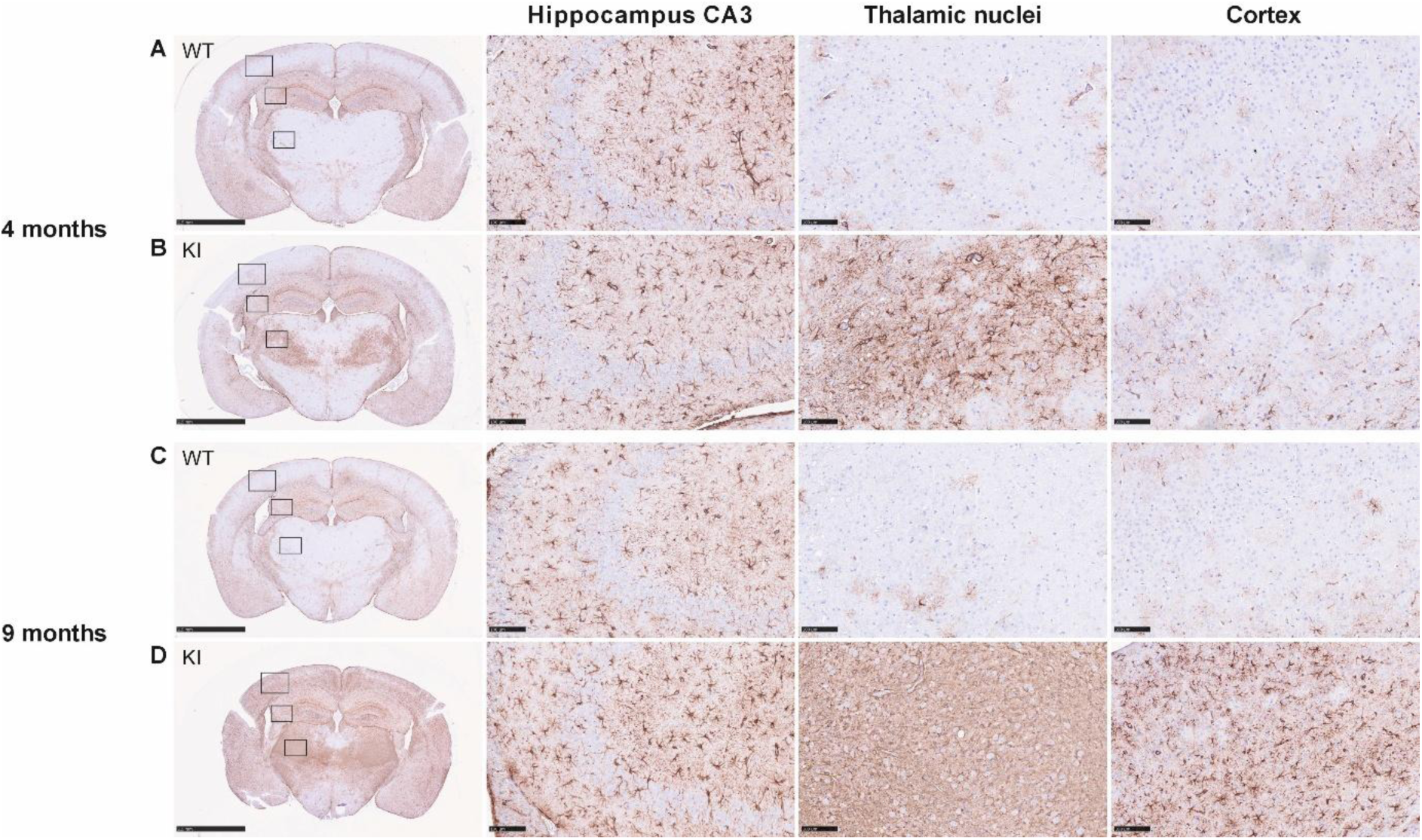
GFAP immunostaining of activated astrocytes indicating neuroinflammation. Astrocyte activation was increased in the thalamic VP nuclei and somatosensory cortex of the Cln8^R24G^ KI males at 4 months of age (B). The activation increased markedly with age (D). Similar increase was not detected in the thalamic VP nuclei and somatosensory cortex of littermate controls (A and C). Activated astrocytes were evident in the hippocampus of both the Cln8^R24G^ KI mice and the littermate controls at 4 and 9 months, at comparable levels (AD). Representative images are selected from male mice (4 months: WT *n* = 4, KI *n* = 4; 9 months: WT *n* = 4, KI *n* = 4). Scale bars: overview, 2.5 mm; hippocampus, thalamic nuclei and cortex, 100 µm.

**Figure 10.**
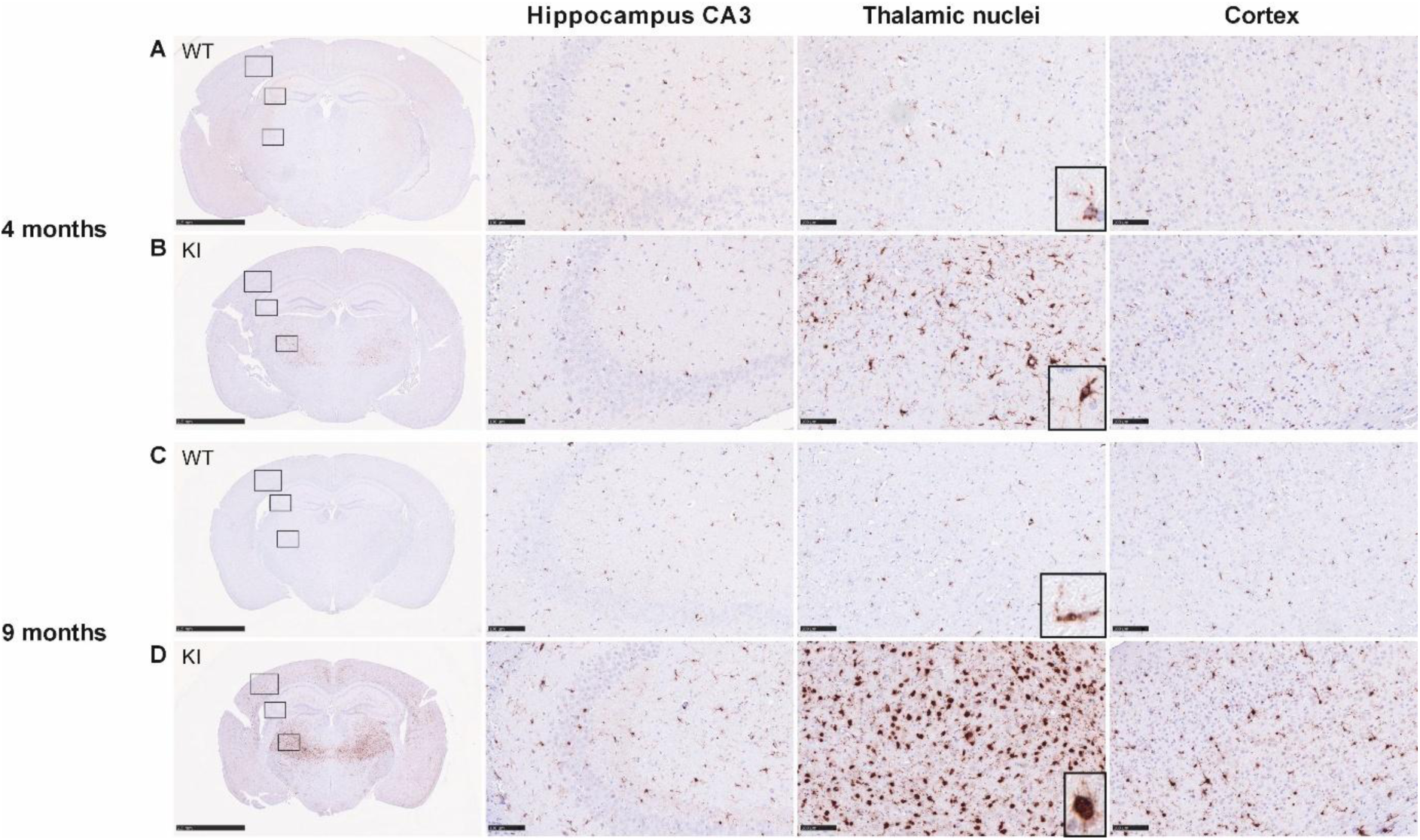
CD68 immunostaining of activated microglia in the hippocampal region. CD68 -positive (CD68^+^) cells were markedly increased in the thalamic VP nuclei of the Cln8^R24G^ KI mice at 4 and 9 months of age (B and D). Increased numbers were also observed in the somatosensory cortex, though less prominent than in the thalamic nuclei. Morphologic changes in the CD68^+^cells with age were observed in the Cln8^R24G^ KI mice (B and D, close-up images): the cells changed from small somata with multiple long processes at 4 months to intensely stained, thick somata with fewer, shorter processes at 9 months. Such changes were absent in the littermate controls (A and C). No differences were evident in the hippocampus between the Cln8^R24G^ KI mice and the littermate controls. Representative images are selected from male mice. Sample sizes: 4 months: WT *n* = 4, KI *n* = 4; 9 months: WT *n* = 4, KI *n* = 4). Scale bars: overview, 2.5 mm; hippocampus, thalamic nuclei and cortex, 100 µm.

Activated microglia were especially evident in layer 5 of the motor cortex (Supplementary Figure 5D) and in the cerebellar nuclei (Figure 11D). No clear differences in astrocyte or microglial activation were observed in the hippocampus between the Cln8^R24G^ KI mice and littermate controls (Figure 10 A-D). The anterior commissure showed a marked increase in activated microglia (Supplementary Figure 5D). Similar findings were noted in the founder line B brains (Supplementary Figure 6).

**Figure 11.**
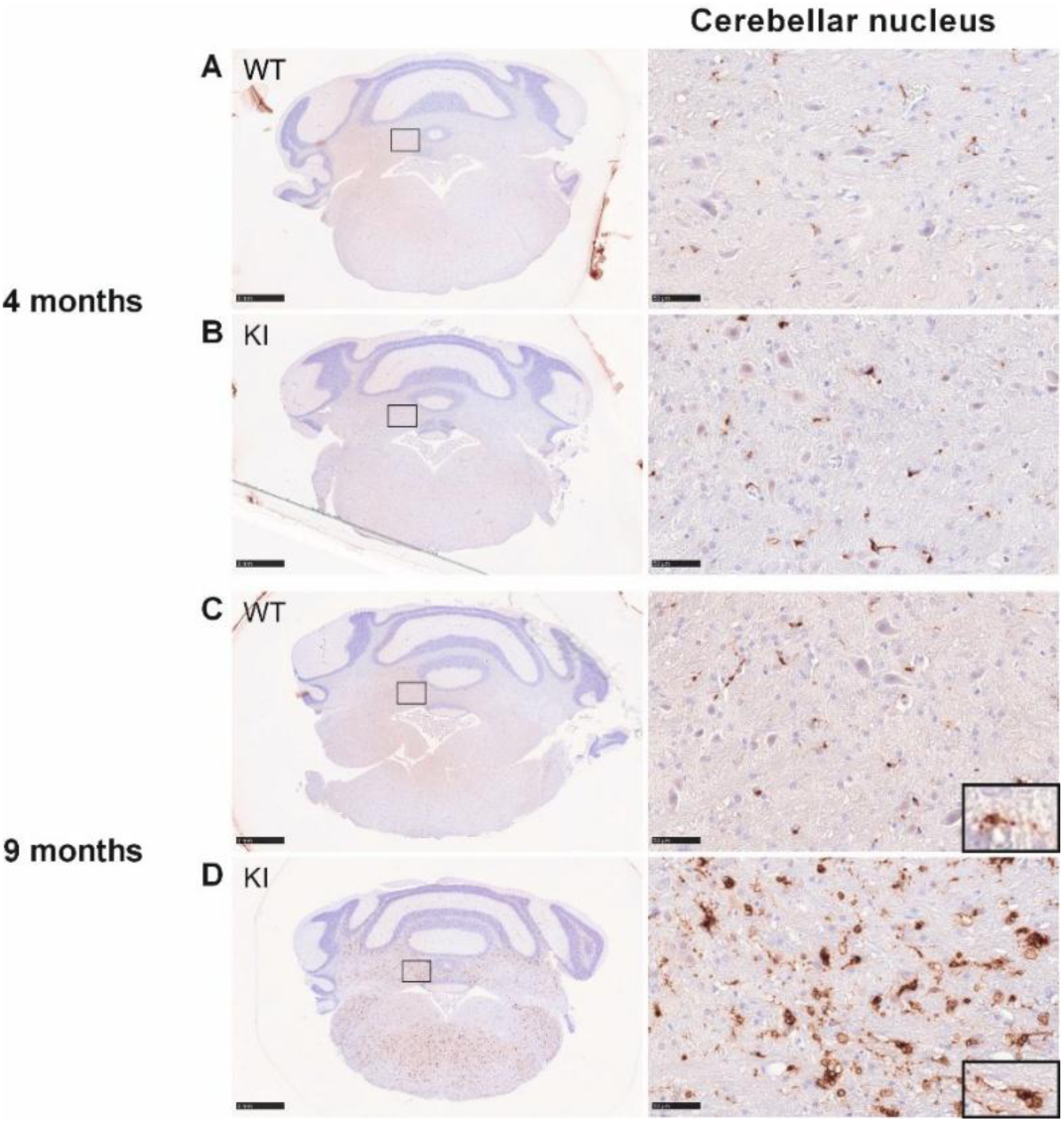
CD68 immunostaining of activated microglia in the cerebellum. The CD68^+^ cells were clearly increased in the medial cerebellar nucleus of the Cln8^R24G^ KI mice at 9 months (D). Many appeared vacuolated (close-up) in the Cln8^R24G^ KI mice (D) but not in the littermate controls (C). No differences were observed at 4 months (A and B). Representative images are selected from male mice. Sample sizes: 4 months: WT *n* = 2, KI *n* = 2; 9 months: WT *n* = 3, KI *n* = 3). Scale bars: overview, 1 mm; cerebellar nucleus, 100 µm.

As the mice aged, the morphology of the activated microglial cells changed. In the 9- versus 4-month-old mice, microglial cell processes were shorter, and the soma was thicker and stained more intensely for CD68 (Figure 10B and D). Strong vacuolisation of activated microglia was evident in the cerebellar nuclei (Figure 11D) but not in the other brain regions. Such vacuolisation was absent in the littermate controls (Figure 11A and C). Low levels of astrocyte and microglial activation were observed in the littermate controls at both ages, but the magnitude was far less than in the Cln8^R24G^ KI mice (A and C in Figures 9, 10, 11). Images for female mice are presented in Supplementary Figure 7.

Brain atrophy is a common feature of Northern epilepsy, and the accumulation of ceroid lipofuscin in brain regions has been correlated with progressive neuronal loss. The amount of NeuN-positive (NeuN^+^) neurons in the cortex (somatosensory barrel field) and thalamic nuclei (VPN/VPL) was not changed in the Cln8^R24G^ KI mice at 4 months, but at 9 months, significant NeuN^+^ cell loss was observed in the thalamic VP nuclei (Figure 12A and B, Supplementary Figure 8A-D, Supplementary Figure 9A). An increase in the NeuN -negative (NeuN^-^) cells was observed in the thalamic nuclei and cortex of Cln8^R24G^ KI mice at 9 months (Figure 12C, Supplementary Figure 9B). The density of NeuN nuclei was not quantified in the hippocampus due to the high cell density, which made it difficult to detect individual cells.

**Figure 12.**
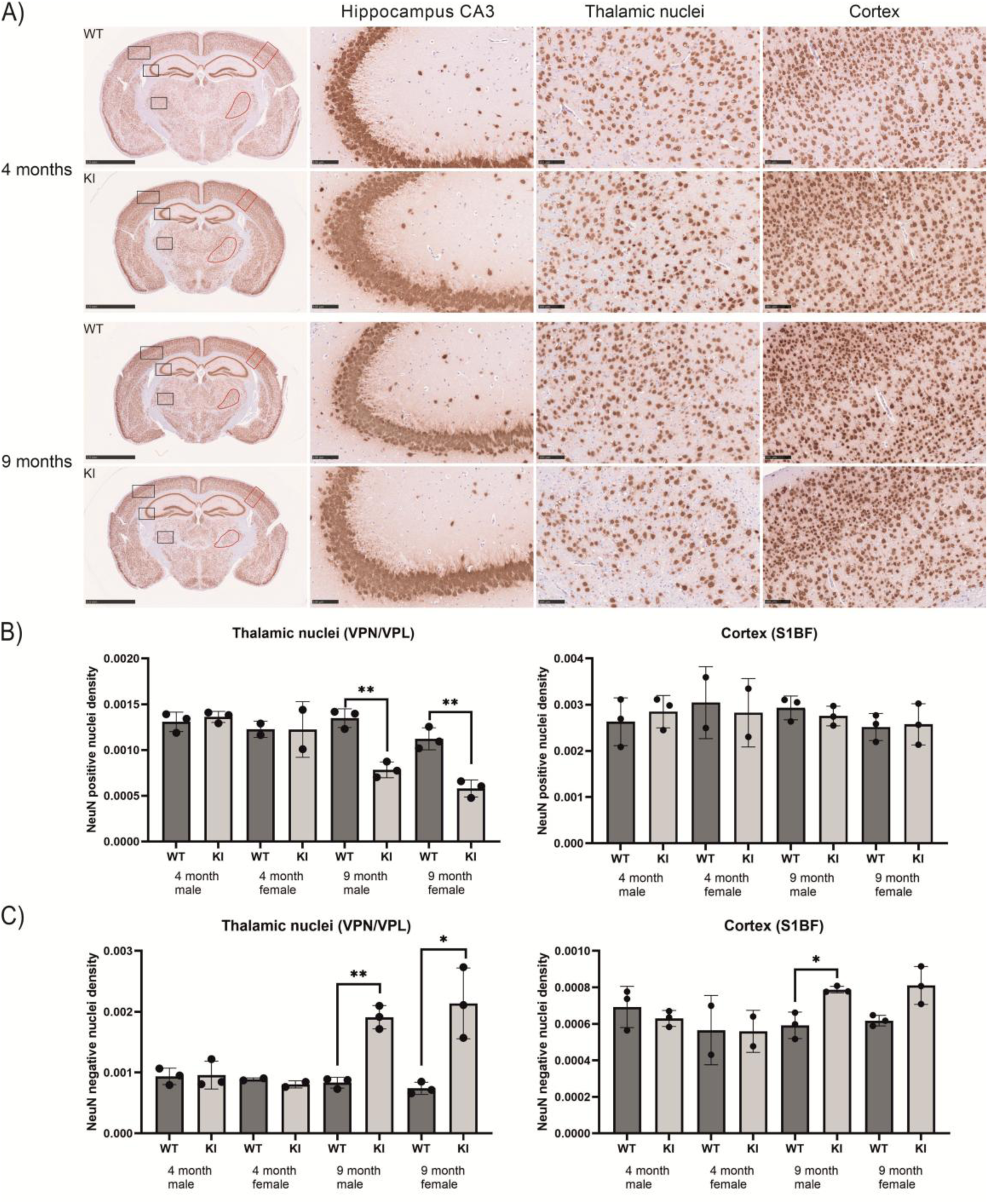
Immunohistochemical detection and quantification of neuronal loss in Cln8^R24G^ KI mice. A) Immunostaining using an antibody against the neuronal marker NeuN showed a significantly decreased neuronal number in the thalamic nuclei of the Cln8^R24G^ KI mice at 9 months of age. No clear differences in the neuronal number were observed in the hippocampus and cortex compared to the littermate controls at either age points. Scale bars: overview, 2.5 mm; hippocampus, thalamic nuclei and cortex, 100 µm. B) Quantification of the NeuN-positive (NeuN^+^) nuclei and C) NeuN -negative (NeuN^-^) nuclei in the thalamic nuclei (VPN/VPL) and somatosensory cortex barrel field 1 (S1BF) of the right brain hemisphere was performed using Visiopharm software. At 9 months, both the male and female Cln8^R24G^ KI mice showed a significant decrease in the NeuN^+^ nuclei density and a corresponding increase in the NeuN^-^ nuclei density in the thalamic nuclei. In the cortex, the NeuN^-^ nuclei density increased in the 9-month-old Cln8^R24G^ KI mice but reached statistical significance only in male mice. No significant differences in the NeuN^+^ nuclei density were observed in the somatosensory cortex at either age. \**P* < 0.05, \*\**P* < 0.01, \*\*\**P* < 0.001. Sample sizes: 4month males, WT *n* = 3, KI *n* = 3; 4-month females, WT *n* = 2, KI *n* = 2; 9-month males, WT *n* = 3, KI *n* = 3; 9-month females, WT *n* = 3, KI *n* = 3. Representative images are selected from male mice. Images from female mice as well as the quantification of NeuN^+^ and NeuN^-^ are presented in Supplementary Figures 8 and 9 respectively.

### Cln8^R24G^ KI mice develop epileptic seizures

Video EEGs were recorded to assess possible epileptic activity in four Cln8^R24G^ KI and two littermate controls at 6−7 months and 8−9 months (Table 1). The analysis focused on cortical and hippocampal epileptiform discharges (EDs). The cortical and hippocampal EDs mainly occurred during the sleep phases in the Cln8^R24G^ KI mice. Therefore, the ED count was normalised to the total sleep time. Cortical EDs were mostly bilateral and followed by brief (∼ 1 s) amplitude increases in the EMG (Figure 13A), characteristic of myoclonus. EDs in the hippocampus had high amplitudes (+/- 2 mV) and were followed by changes in the background field potential (Figure 13B). Wildtype and two Cln8^R24G^ KI mice showed no EDs at 6−7 months, but the mutants showed occasional cortical and hippocampal EDs at 8−9 months (Table 1). The other two Cln8^R24G^ KI mice already exhibited a substantial number of cortical and hippocampal EDs at 6−7 months. The number of cortical spikes in these mice did not increase with age, but hippocampal EDs became more frequent (> 90 per hour = 1.5 EDs per minute). One of the mutant mice exhibited a generalised tonic−clonic seizure during the final 3-h recording session (Figure 13C), followed by status epilepticus (Figure 13D), before its brain was perfused for histology.

**Figure 13.**
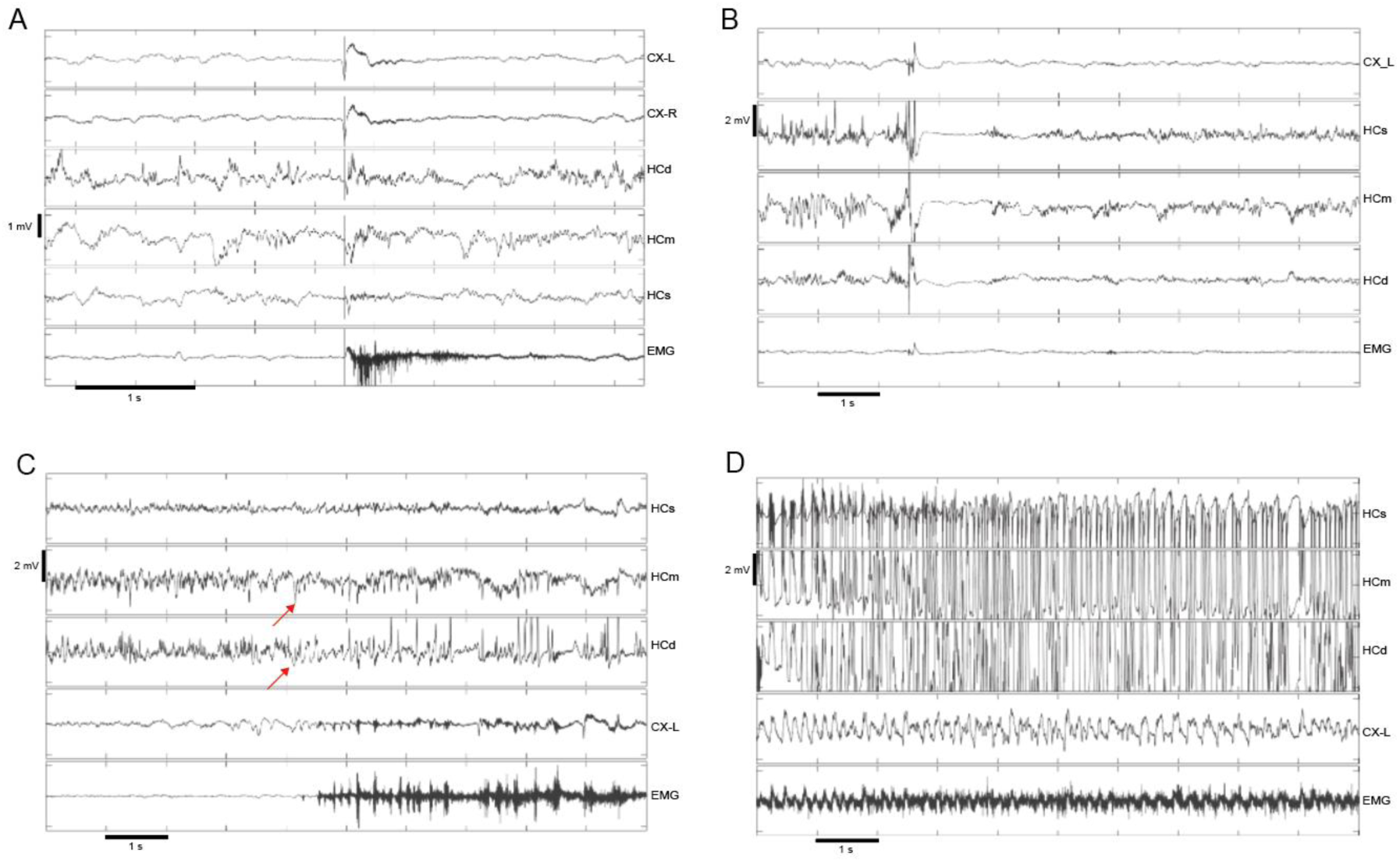
Epileptiform EEG findings in the KI-4 mouse. (A) Bilateral cortical ED with myoclonus. (B) Massive ED originating in the hippocampus, followed by 1.5s suppression of local electric activity. (C) Onset of a generalised seizure. The red arrows indicate the seizure origin in the deep hippocampal channels (dentate gyrus). The seizure was almost immediately followed by increased muscle tone in the EMG indicated generalized seizure starting with a tonic phase. (D) Fully developed seizure characterised by high-amplitude rhythmic spikes and waves across all channels. The rhythmic EMG changes corresponded to the clonic phase. Channel abbreviations: cortical screw electrode, CX; hippocampal triple wire electrode, HC (s = superficial, m = middle, d = deep); electromyograph, EMG.

**Table 1.** Densities of cortical and hippocampal epileptiform discharges (EDs) in Cln8^R24G^ KI and WT mice. The ED densities were recorded in the same animals at 6−7 and at 8−9 months of age. HC = hippocampal EDs; CX = cortical EDs. The values represent the number of EDs per hour of sleep.

| Mouse ID | HC (6–7 mo) | HC (8–9 mo) | CX (6–7 mo) | CX (8–9 mo) |
| --- | --- | --- | --- | --- |
| WT-1 | 0 | 0.5 | 0 | 3.6 |
| WT-2 | 0 | 0 | 0 | 0.9 |
| KI -1 | 0 | 1.8 | 0 | 2.7 |
| KI -2 | 0 | 1.1 | 0 | 3.2 |
| KI -3 | 5.9 | 91.2 | 2.0 | 3.0 |
| KI -4 | 14.7 | 91.8 | 85.3 | 15.3 |

*Note: Mouse KI-4 experienced a generalised seizure followed by status epilepticus at 8.5 months.*

We used a FosB antibody to detect both full-length FosB and its truncated splice variant dFosB. The latter accumulates in the nuclei of chronically active neurons, has a long half-life and thus reflects mean neuronal activity over the preceding 1-3 weeks (Patterson et al., 2016). We examined potential correlations between neuronal hyperactivity and lipofuscin accumulation, and between epileptic activity in the EEG and FosB expression. To this end, we analysed the brains of two Cln8^R24G^ WT and four Cln8^R24G^ KI mice with successful EEG recordings. All four Cln8^R24G^ KI mice differed from the two WT mice in FosB staining intensity (*p* = 0.02, Mann-Whitney U), most prominently in the dentate gyrus and piriform cortex (Figure 14A). Increased FosB staining was also observed in CA1 and the dorsal cortical convexity, though to a lesser extent. The Cln8^R24G^ KI-4 mouse, which was perfused after showing status epilepticus (Figure 13 and Table 1), exhibited the strongest FosB staining, extending across all hippocampal neuronal layers and the entire cortical mantle (Figure 14A). Intense FosB staining was also evident in the left amygdala. Autofluorescence in the same mouse showed typical increases in hippocampal pyramidal cell layers and thalamic nuclei (Figure 14B). Interestingly, the FosB and autofluorescence staining patterns appeared complementary (Figure 14A and B). For example, in the hippocampus, CA pyramidal cells displayed the strongest autofluorescence, whereas the FosB intensity was highest in the dentate granule cell layer. In the cortex, the piriform cortex exhibited intense FosB staining but low autofluorescence compared with other cortical regions.

**Figure 14.**
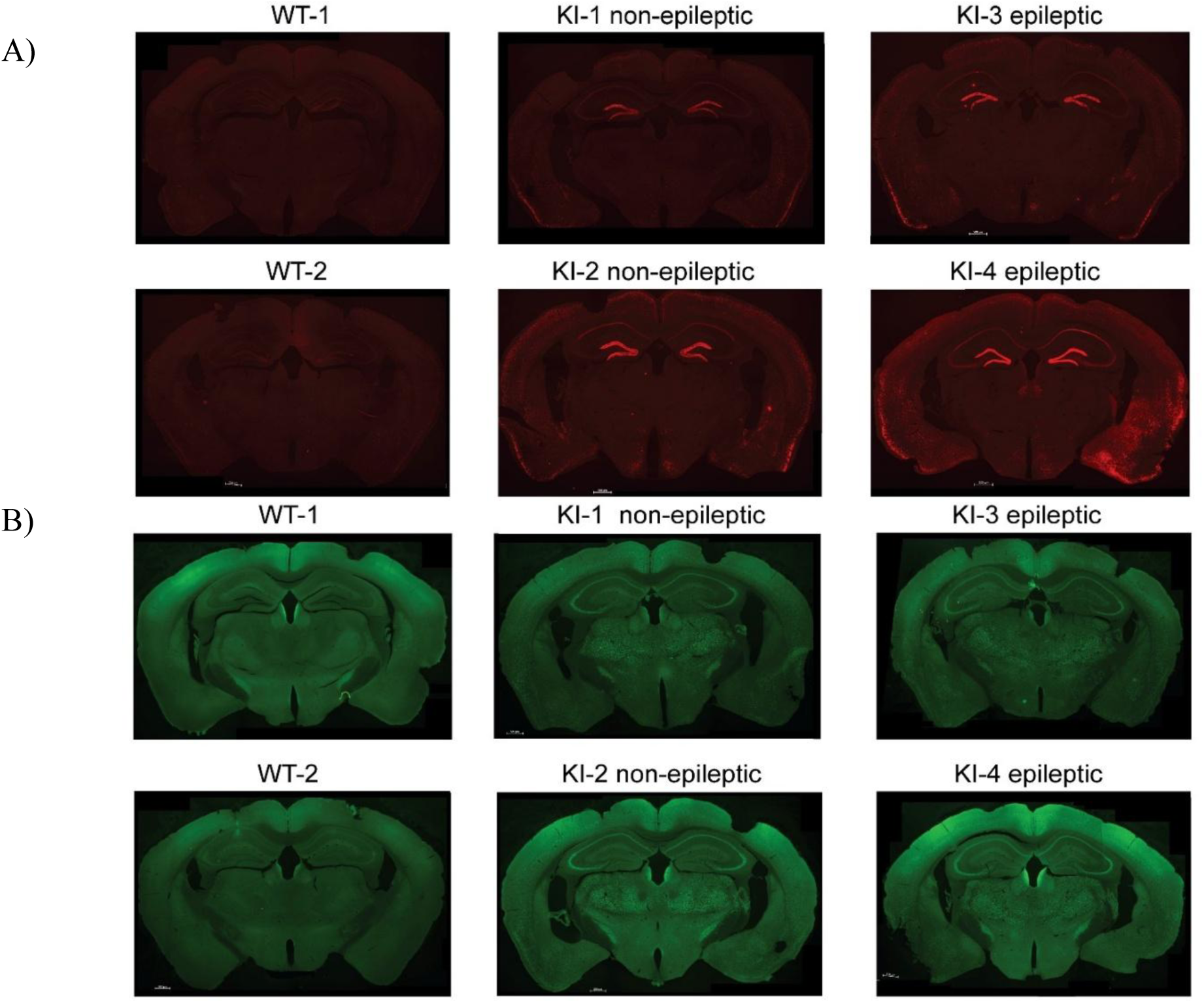
Visualization of the neuronal hyperactivity in Cln8^R24G^ KI mice after EEG. A) FosB staining showed no region that stood out from the background in the two Cln8^R24G^ WT mice, whereas all Cln8^R24G^ KI mice exhibited prominent staining in the dentate granule cell layer and piriform cortex. The only exception was mouse KI-4 (bottom right), which had a seizure and status epilepticus; this mouse displayed intensified FosB staining across all hippocampal fields, and strong staining in the left amygdala. B) Autofluorescence patterns differed clearly between the WT and KI mice but did not show marked differences between the two epileptic and two non-epileptic KI mice.

### Accumulation of autofluorescent storage material and progressive astrogliosis in the retina of Cln8^R24G^ KI mice

In addition to the brain-related findings, we examined whether early pathological signs could be detected in the retina. Granular autofluorescent storage material was present and progressively accumulated in the Cln8^R24G^ KI, but not in the WT mice (Figure 15). The signal was most prominent at the level of the retinal ganglion cells (RGCs), between inner plexiform (IP) and nuclear (IN) layers, and at the outer plexiform layer (OPL).

**Figure 15.**
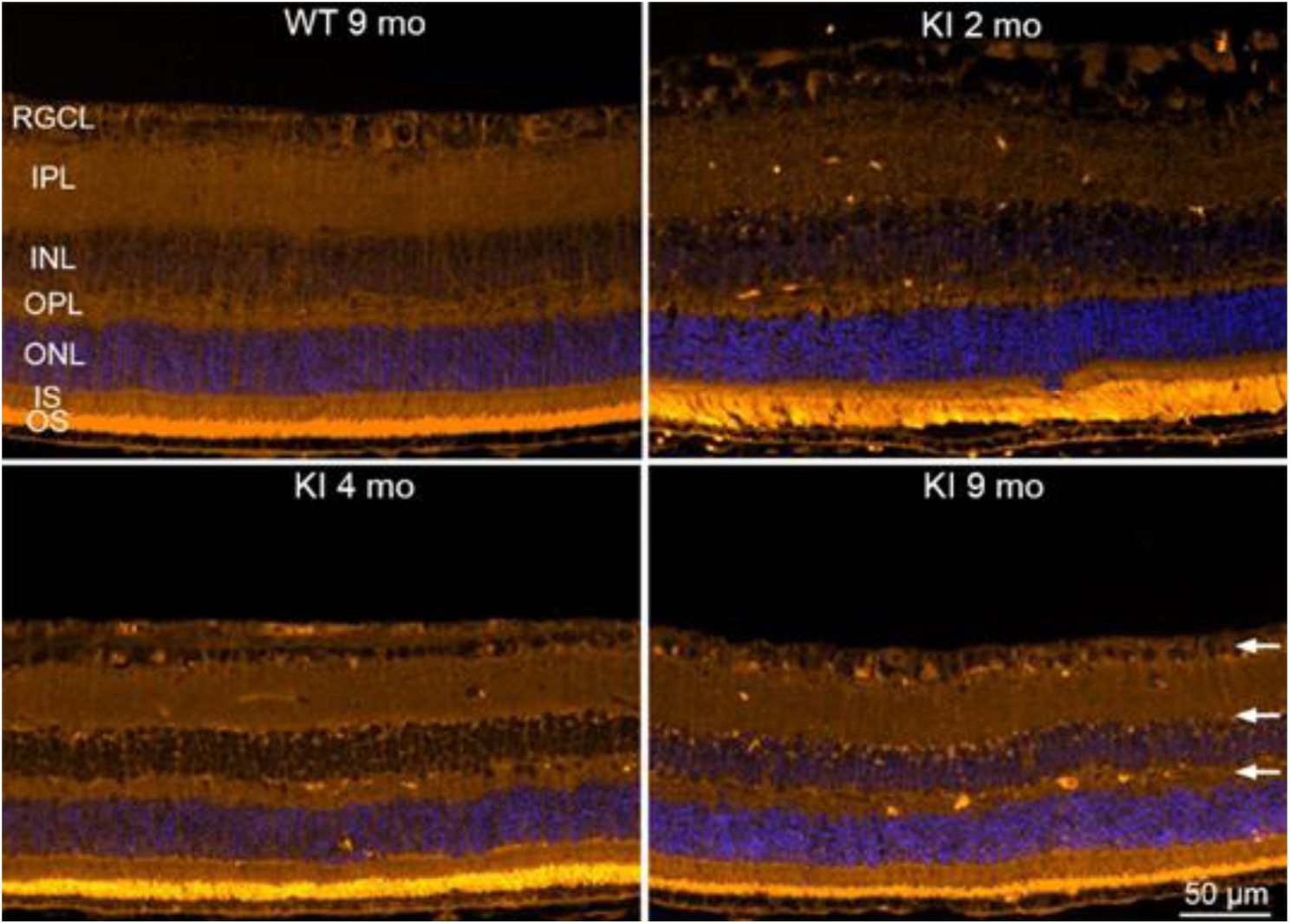
Autofluorescent accumulation in retina. Autofluorescent signal in retina crosssections. The autofluorescence was detectable as early as 2 months in the Cln8^R24G^ KI mice, with progressive accumulation observed in the retinal ganglion cell layer (RGCL), inner nuclear layer (INL, particularly at the INL−IPL junction) and outer plexiform layer (OPL), indicated with arrows. Sample sizes: KI 2 mo, *n* = 3; KI 4 mo, *n* = 10; KI 9 mo, *n* = 3. Similar autofluorescence signal was absent in Cln8^R24G^ WT mouse retinas (*n* = 3). Abbreviations: retinal ganglion cell layer, RGCL; inner plexiform layer, IPL; inner nuclear layer, INL; outer plexiform layer, OPL; outer nuclear layer, ONL; inner segment layer, IS; outer segment layer, OS; WT, wild type; knock-in, KI; mo, month.

GFAP-immunohistochemistry revealed modestly reactive astrogliosis in the 4-month-old Cln8^R24G^ KI mice, which became more pronounced by 9 months (Figure 16). At this later time point, activation of Müller glia was also evident. These retinal changes parallel the progressive increase in GFAP^+^ cells observed in the brain between 4 and 9 months. In addition, retinal degeneration was observed, particularly at the outer nuclear layer where photoreceptor nuclei reside. The M-opsin staining confirmed that the location of GFAP signal and outer nuclear layer thickness observation was at the superior half of the retina.

**Figure 16.**
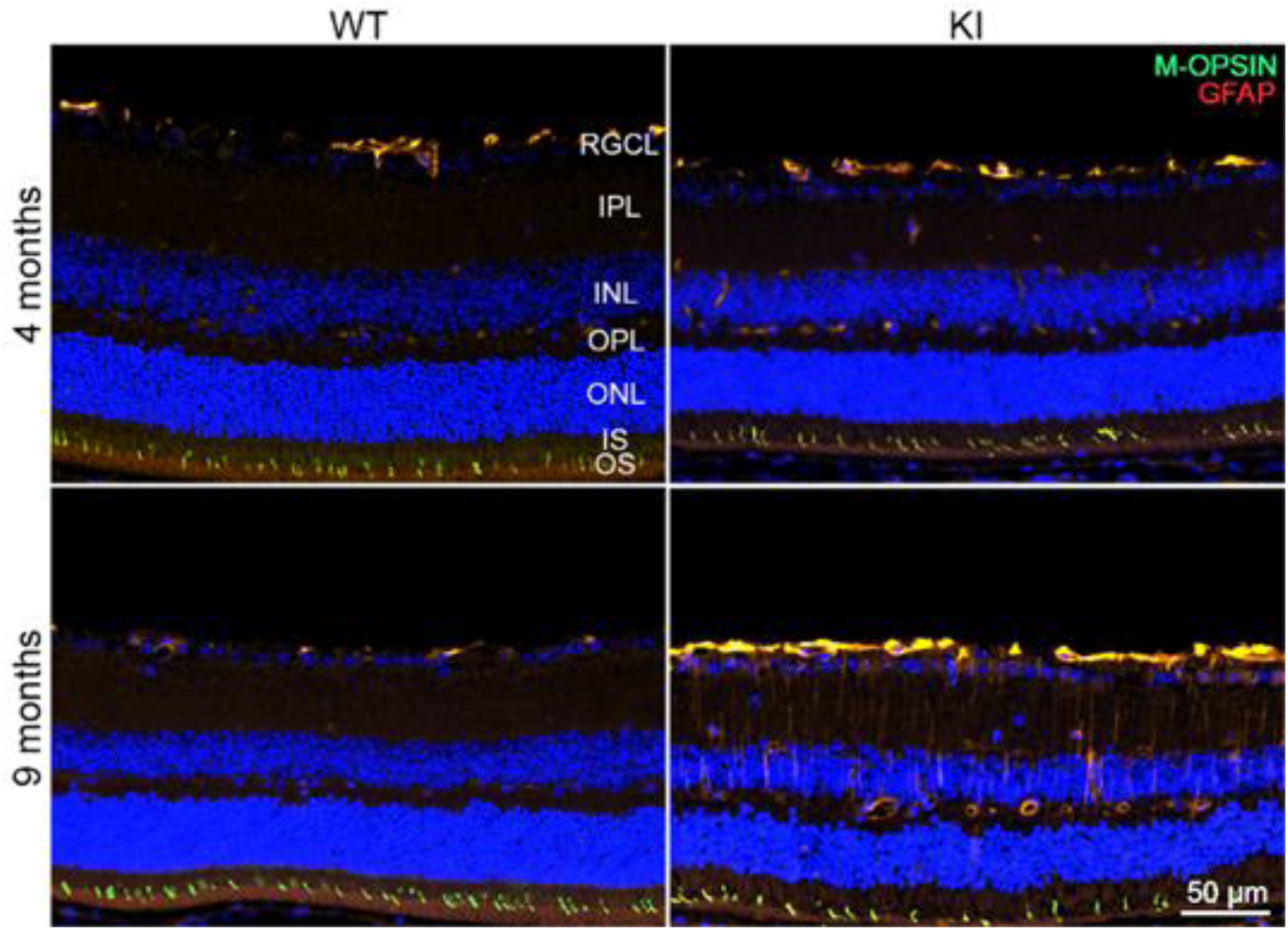
Immunohistochemical staining using anti-GFAP and anti-M-opsin antibodies in the retinas of Cln8^R24G^ mice. GFAP, a widely used gliosis marker, revealed inflammatory activity in the retinal ganglion cell layer (RGCL) of the 4-month-old Cln8^R24G^ KI mice, consistent with astrogliosis. In the 9-month-old Cln8^R24G^ KI mice, the inflammatory response was elevated, accompanied by clear activation of the radially projecting Müller glia. Sample sizes: 4 months, WT *n* = 8, KI *n* = 10; 9 months, WT *n* = 3, KI *n* = 3. Abbreviations: retinal ganglion cell layer, RGCL; inner plexiform layer, IPL; inner nuclear layer, INL; outer plexiform layer, OPL; outer nuclear layer, ONL; inner segment layer, IS; outer segment layer, OS; WT, wild type; knock-in, KI.

## Discussion

Here, we describe the first Cln8^R24G^ mouse model that genocopies the pathogenic variant found in patients with Northern epilepsy. This mouse model recapitulates the clinical features of Northern epilepsy and shows classical histopathological features of Batten disease in both the brain and retina. These include a progressive disease course characterized by retinal degeneration, motor deterioration and epileptic seizures associated with the progressive accumulation of autofluorescent storage material in the retina and brain, together with neuroinflammation and neurodegeneration in the CNS.

Accumulation of autofluorescent ceroid lipofuscin was evident in both the brain and retina from the earliest ages analysed (brain, 4 months; retina, 2 months) and progressively increased with age. The storage material was primarily seen in neurons, stained positive for SCMAS and phospholipids (LFB^+^). Ultrastructurally, it appeared granular and curvilinear-like as in Northern epilepsy patients (Haltia, 2003), differing from lipofuscin that accumulates during normal ageing (Terman & Brunk, 1998). Accumulation of autofluorescent storage material has also been reported in the mnd mouse model, which carries a mutation not found in humans (Bronson et al., 1993). Nonetheless, Cln8 deficiency in mnd mice, Cln8^R24G^ mice and Northern epilepsy patients results in comparable pathological accumulations of ceroid lipofuscin.

The retinal pathology in Cln8^R24G^ KI mice resembled findings in mnd mice but was milder and slowly progressive, more closely mimicking the clinical course of Northern Epilepsy patients, in whom visual impairment is not among the main clinical symptoms. In mnd mouse retinas, autofluorescent storage material accumulates prominently within the RGC layer and inner retina at early postnatal stages (Seigel et al., 2005). By one month of age, robust ER-stress responses and apoptosis indicate severe retinal degeneration (Galizzi et al., 2011; Guarneri et al., 2004; Seigel et al., 2005), leading to nearly complete photoreceptor loss by 8 months, particularly in female mnd mice (Guarneri et al., 2004). In contrast, Cln8^R24G^ KI mice showed preserved outer nuclear layer thickness and photoreceptor inner/outer segments at age 2 months, with only modest thinning by 4 months. The inner retina retained normal lamination without major macrostructural disruption up to at least 9 months. GFAP immunoreactivity emerged by 4 months and became more pronounced by 9 months, consistent with progressive gliosis observed in other CNS regions such as the thalamus. Despite degeneration affecting primarily the outer retina (photoreceptors), granular autofluorescence was detected mainly in the RGC layer and at the border between the IP and IN layers. Larger deposits occurred occasionally in the synaptic plexiform layers across all ages examined (2, 4 and 9 months). Taken together, these findings and previous reports suggest that lysosomal storage pathology arises early in the inner retina across different Cln8 models but does not strictly correlate with the sites of structural degeneration. Interestingly, the relatively slow progression of retinal pathology in Cln8^R24G^ KI mice more closely resembles that seen in *Cln8* full knockout mice (Salpeter et al., 2021), than the severe retinal degeneration observed in mnd mice.

In the brain, ceroid lipofuscin most prominently accumulated in the hippocampus (CA3 sector), cerebellum (Purkinje cells and medial cerebellar nucleus), somatosensory and motor cortex, and thalamic VP nuclei interconnected with somatosensory and motor cortex. This distribution resembles the pattern reported in mnd mice (Kuronen et al., 2012; Pardo et al., 1994), although the accumulation of autofluorescence in the cerebellum has been less extensively studied (Galizzi et al., 2011). In CLN8 patients, the accumulation of intraneuronal storage material has been described mainly in the cortex (third layer) and hippocampus (CA2, CA3, and CA4) (Herva et al., 2000; Tyynelä et al., 2004). Interestingly, other layers of the isocortex, CA1 and cerebellar cortex appear only minimally affected in patients. The basal ganglia, thalamus, brainstem and spinal cord have been reported to show slight to moderate intraneuronal storage (Haltia, 2003). Differences between species may explain some inconsistencies in these findings, but they could also reflect variation in disease stages at the time of sampling and the limited availability of patient-derived samples.

Neuroinflammation, assessed by increased activation of astrocytes and microglial cells, has been reported to precede neuronal loss in mnd mice and in CLN8 patients (Herva et al., 2000; Kuronen et al., 2012; Lauronen et al., 2001; Tyynelä et al., 1993). In the Cln8^R24G^ KI mice, astrogliosis and microgliosis were evident at 4 months of age and progressively increased between 4 and 9 months in the VP nucleus of the thalamus, with more moderate changes in the cortex. Microglial cell morphology also changed over time: in the thalamus, processes became shorter, and the soma became thick and CD68^+^-intensive at 9 months, features characteristic of microglial activation (Helmut et al., 2011; Hendrickx et al., 2017). At 9 months, a significant decrease in NeuN^+^ neurons was observed in the thalamic VP nuclei, but not in the cortex, suggesting a correlation between neuroinflammation and neurodegeneration in the thalamic VP nucleus. No clear increase in astrocyte and microglial activation was detected in the hippocampus of the Cln8^R24G^ KI mice despite prominent accumulation of autofluorescent material in this region. In contrast, the anterior commissure showed marked microglia activation but little autofluorescence. These findings indicate that accumulation of storage material does not necessarily correlate with the activation of neuroinflammation and neurodegeneration, except in specific regions such as the thalamic nuclei. Similar to mnd mice (Kuronen et al., 2012), the thalamic VP nuclei were the most prominently affected brain regions in the Cln8^R24G^ mice, showing the highest levels of microglia activation and neuronal loss at later stages. Interestingly, altered signal intensity in the thalamus is one of the earliest changes detected by magnetic resonance imaging in patients with late-infantile variant CLN8 disease, suggesting particular vulnerability of the thalamus at early disease stages (Autti et al., 2007; Lauronen et al., 2001; Topçu et al., 2004). In the autopsy material from Northern epilepsy patients, glial activation appears to coincide with neuron loss in hippocampal subfields CA2 and CA3, other brain regions have not been examined in detail (Herva et al., 2000; Tyynelä et al., 2004).

Importantly, the Cln8^R24G^ KI mice recapitulated the main neurobehavioural features of Northern epilepsy, including motor deterioration and epileptic seizures. Motor impairment affecting balance and gait was evident from 7 months onwards. Combining this late-onset symptom with our histopathological findings, it can be hypothesized that motor deterioration is connected with neurodegeneration in the thalamic VP nucleus, an important node in extrapyramical loops, and in other brain regions affected by progressive autofluorescence accumulation, especially cerebellar Purkinje cells and the medial cerebellar nucleus. Motor symptoms have also been described in mnd mice with the C57B6 background, but they appear more severe and typically occur earlier (at around 6 months of age) (Johnson et al., 2021). Spontaneous epileptic seizures resembling those in Northern epilepsy patients (generalised tonic-clonic seizures and myoclonic jerks) were detected and recorded. Increased FosB staining intensity, indicative of neuronal hyperactivity in the hippocampus, correlated well with the seizure activity, suggesting the involvement of hippocampal CA1−CA3 pyramidal cells and dentate granule cells in the epileptogenesis of these mice. In contrast, to our knowledge, no spontaneous epileptic seizures have been reported in mnd mice. Depending on the genetic background, motor impairment in mnd mice progresses to severe spastic paralysis and premature death between 7 and 12 months of age (Messer et al., 1995). The mnd mouse model originates from a naturally occurring frameshift variant (c.267–268insC) predicted to introduce a premature stop -codon after 27 amino acids and truncation of the Cln8 protein (Ranta et al., 1999). Thus, the key difference between the phenotypes of these two mutant mouse models may reflect the distinct nature of their underlying pathogenic variants, with the amino acid substitution p.R24G in Cln8 potentially playing a significant role in the induction of epileptic seizure activity.

Currently, the NCL resource lists 57 putatively pathogenic variants reported in CLN8 patients (http://www.ucl.ac.uk/ncl/). *CLN8* encodes a ubiquitously expressed protein of 286 amino acids (Ranta et al., 1999). It is 33kDa in size and is predicted to contain seven transmembrane domains and C-terminal ER-export and retrieval signals (di Ronza et al., 2018; Sheokand et al., 2025). Sequence homology links CLN8 to the large eukaryotic protein family of TLC-domain homologs (Winter & Ponting, 2002; Sheokand et al., 2025). The pathogenic p.R24G variant is located at the border of the first predicted transmembrane domain and outside the TLC domain, which is composed of residues p.62-262 (Kousi et al., 2012; Winter & Pontig, 2002). The hydrophobicity prediction was not significantly changed when p.Arg24 was replaced by glycine (Ranta et al., 1999), suggesting that the substitution may cause only subtle structural and functional changes in the CLN8 protein. Due to the lack of a specific antibody, we were unable to quantify the mutant Cln8, but the quantitative RT-PCR results demonstrated that *Cln8* expression remained unaltered in the Cln8^R24G^ KI mice. Compared with the mnd mouse model, in which significant truncation of the Cln8 protein is expected, it is surprising how similar the histopathological phenotypes appear. On the other hand, the differences between phenotypes may reflect the presence of the TLC domain, which is presumably completely lost in the mnd mice but remains unaltered in our Cln8^R24G^ mouse line.

Despite the well-characterised disease phenotype and genetic aetiology of Northern epilepsy, no curative treatment is currently available. The development of effective therapies remains as an unmet need and a major challenge in ongoing research. Therefore, a deeper understanding of the underlying molecular mechanisms is critical. Mouse models have traditionally played a major role in such investigations, as many histological and biochemical studies in the most affected organ, the central nervous system, cannot be performed in humans. Here, we describe a novel Cln8^R24G^ mouse model of Northern epilepsy. Our data demonstrate significant similarities to the mnd mouse model, which has been widely used to study CLN8 -related diseases, thereby validating this novel mutant mouse line. Moreover, our findings closely align with previously reported observations in CLN8 patients. Notably, the novel Cln8^R24G^ mouse model recapitulates spontaneous epileptic seizures, one of the main clinical features of Northern epilepsy, which have not been reported to occur in the mnd mouse model. Altogether, we hypothesize that our mouse model will open new possibilities for developing and testing targeted treatment options for Northern epilepsy and, more broadly, for early-onset neurodegenerative disorders associated with epileptic seizures.

## Acknowledgements

This work was funded by Jane ja Aatos Erkon Säätiö (Jane and Aatos Erkko Foundation; P.S., S.K., R.H., FinnDisMice consortium); Lastentautien Tutkimussäätiö (Foundation for Pediatric Research, Finland; J.U., R.H.); the Research Council of Finland (profiling programme, 311934, R.H.; 346295, H.L.; 331436, J.U.; 348906, S.K.); the University of Oulu (R.H.); the University of Oulu Graduate School (J.M-N.); the NCL Foundation (J.M-N.; R.H.); Competitive State Funding for Health Research for the Wellbeing Services County of North Ostrobothnia, Finland (J.U.); Emil Aaltonen Foundation (Emil Aaltosen Säätiö; H.L.); Sigrid Jusélius Foundation (Sigrid Juséliuksen Säätiö; L.E.; H.L.); the FEBS Excellence Award 2024 (H.L.); the University of Helsinki (S.K.) and the European Union’s Horizon Europe research and innovation programme under grant agreement No 101131669 (INFRAPLUS; R.H.; A.H.; M.H.S.).

We thank Dr. Herman van der Putten (NCL Foundation) and Prof. Anna-Elina Lehesjoki (Folkhälsan Research Center, University of Helsinki) for their support and expert advice. The following Biocenter Oulu core facilities, supported by the University of Oulu and Biocenter Finland: Transgenic and Tissue Phenotyping, Electron Microscopy Core Facility and Sequencing Center, and The Laboratory Animal Centre at the University of Oulu, are acknowledged for their expert services, resources and advice. The Biocenter Kuopio Phenotyping Center and Laboratory Animal Center at the University of Eastern Finland are acknowledged for their technical support and service. The authors thank Saga Kotkaranta and Pirjo Keränen for their expert assistance in our laboratory experiments.

Some authors of this publication are members of the European Reference Network on Rare Neurological Diseases (ERN-RND), Rare and Complex Epilepsies (EpiCARE), Neuromuscular Diseases (ERN-EURO-NMD) and Rare Congenital Malformations and Rare Intellectual Disability (ERN-ITHACA).

