## Supplementary material for "Novel Cln8 p.R24G mouse line replicates major clinical features of Northern epilepsy"

### Supplementary Data

#### Supplementary Table 1. Primer sequences.

| Target | Primer sequence (5'-3') | Product size (bp) |
|---|---|---|
| *Rpl13a* (Ribosomal protein L13a) | Forward: GAGGTCGGGTGGAAGTACCA<br>Reverse: TGCATCTTGGCCTTTTCCTT | 71 |
| *Hprt1* (Hypoxanthine phosphoribosyltransferase 1) | Forward: CATCATCGCTAATCACGACGC<br>Reverse: CTTCCTCCTCAGACCGCTTT | 85 |
| *Pgk1* (Phosphoglycerate kinase 1) | Forward: CTGACTTTGGACAAGCTGGACG<br>Reverse: GCAGCCTTGATCCTTTGGTTG | 110 |
| *Cln8* (genotyping and sequencing) | Forward: GCCTGAGCGTCTCGGATT<br>Reverse: CTCACCTTCAGGAGCATCCA | 891 |
| *Cln8* (cDNA sequencing) | Forward 1: TGGTGATTTCTCCGGTGC<br>Forward 2: ACCTGTCCAACCTGTTCTT<br>Reverse: AGGCCCCAGCAATACTTCA | |
| *Cln8* (cDNA amplification) | Forward: TGGTGATTTCTCCGGTGC<br>Reverse: AGGCCCCAGCAATACTTCA | 1392 |
| *Cln8* (qPCR) | Forward: CTAGCCATGACCACGTTGCT<br>Reverse: ATTAGCCACTGGTTGGCCTT | 116 |

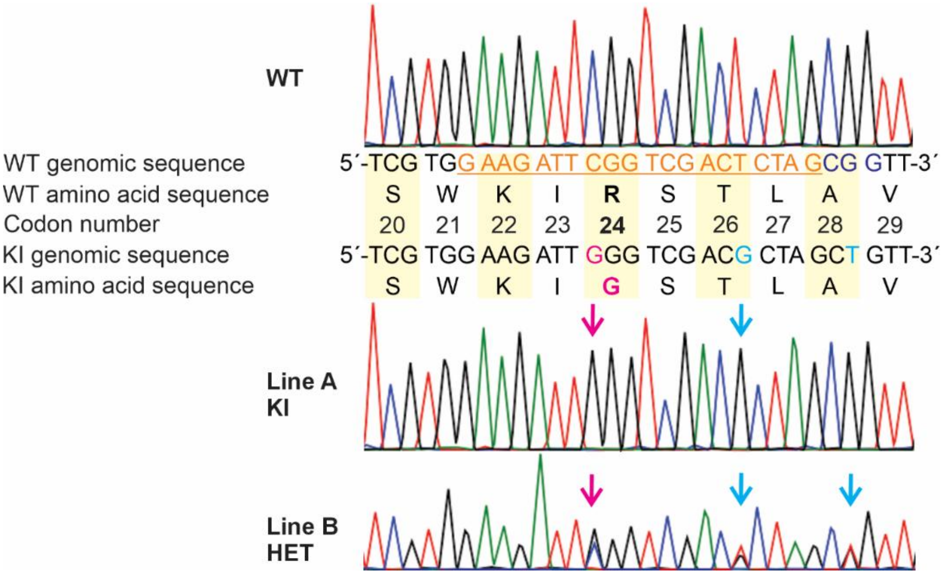


**Supplementary Figure 1**. **Sanger sequencing of heterozygous (HET) $Cln8^{R24G}$ line B mice.** Line B carries the same CGG>GGG modification at codon 24 (ENSMUST00000027554, from the mouse genome assembly GRCm38) and the silent mutation at codon 26 (ACT>AGC, *Nhe*I restriction site) as line A. In addition, line B contains a silent mutation at codon 28 (GCG>GCT) to eliminate the existing PAM site. In the WT genomic sequence, the guide sequence is underlined and highlighted in orange, and the PAM sequence is highlighted in blue.

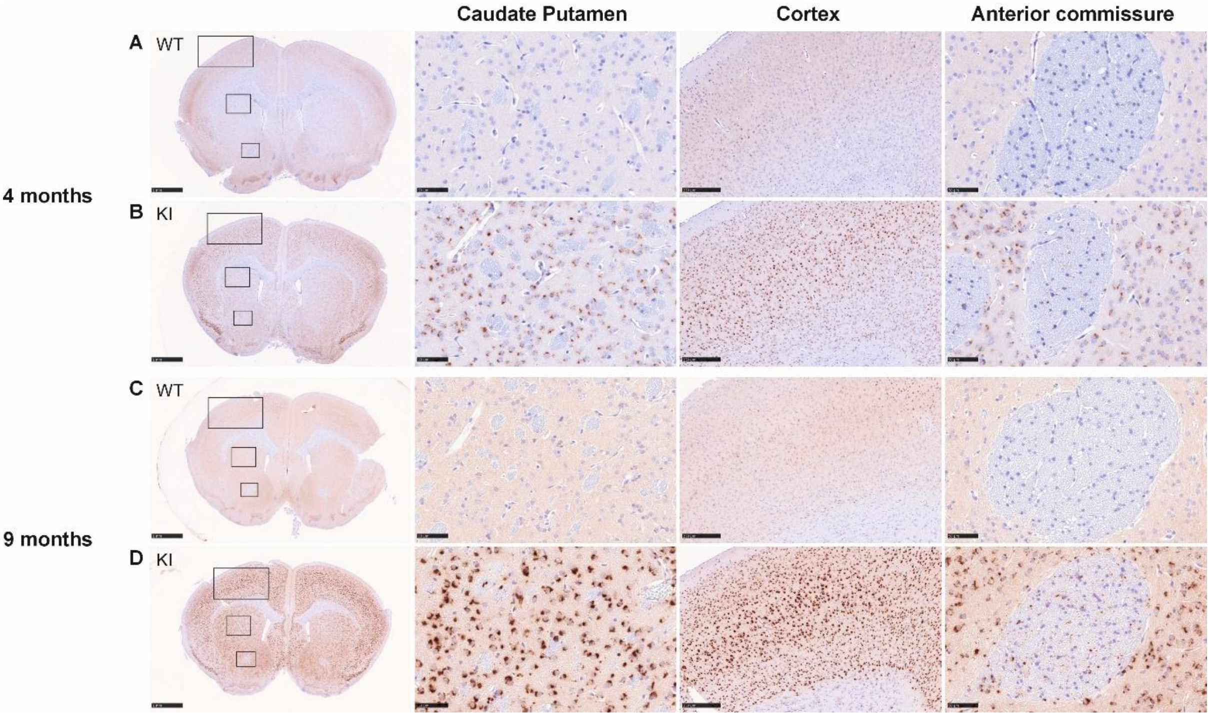


**Supplementary Figure 2. Immunohistochemical SCMAS staining of striatum region in Cln8$^{R24G}$ A-line males.** Immunohistochemical staining in the striatum region showed prominent accumulation of SCMAS in the motor cortex and caudate putamen in the Cln8$^{R24G}$ KI mice (B) but not in the littermate controls at 4 months of age (A). The accumulation increased progressively with age (D). The anterior commissure in the Cln8$^{R24G}$ KI mice showed mild accumulation (B, D) while no SCMAS staining signal was evident in the anterior commissure in the littermate controls (A, C). Representative images are selected from male mice (4 months: WT $n$ = 2, KI $n$ = 2; 9 months: WT $n$ = 3, KI $n$ = 3). Scale bars: overview, 1 mm; caudate putamen and anterior commissure, 50 µm; cortex, 250 µm.

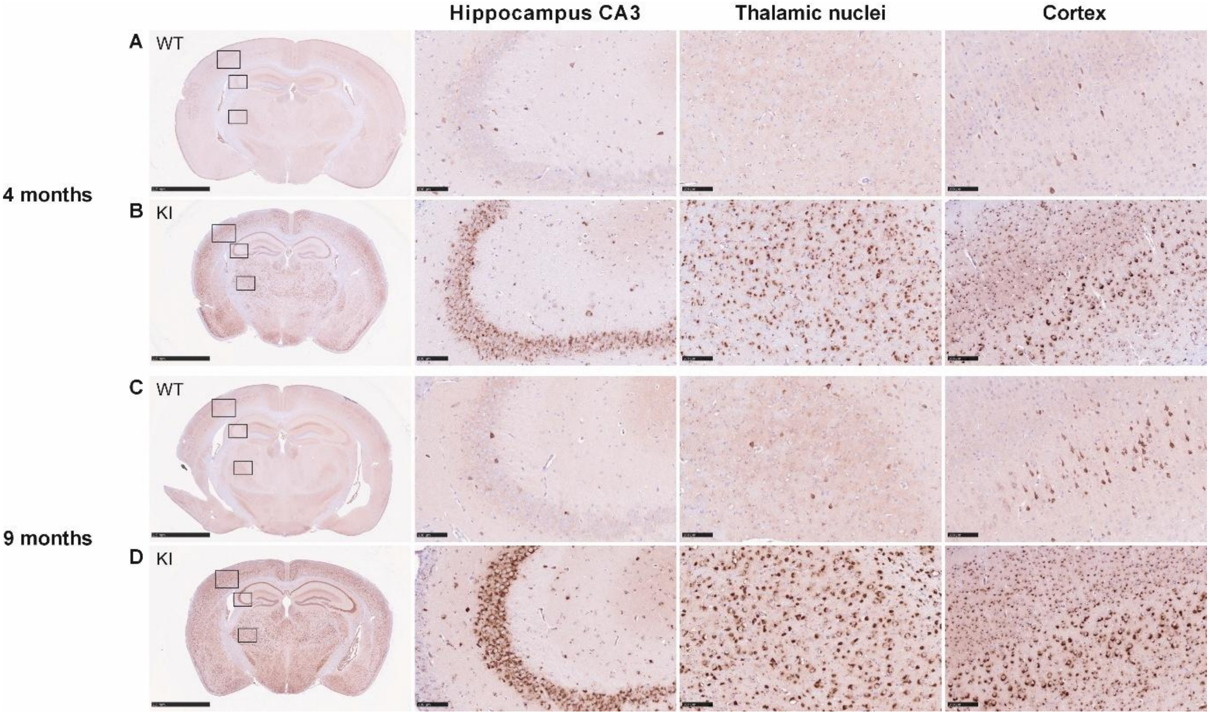


**Supplementary Figure 3. Immunohistochemical SCMAS staining of the hippocampal region of the Cln8$^{R24G}$ A-line females to visualise the accumulation of storage material.** The accumulation was prominent in the hippocampus, VPN/VPL thalamic nuclei and the cortex of the Cln8$^{R24G}$ KI mice (B) but not of the littermate controls from 4 months of age (A). The accumulation was progressive and increased at 9 months in Cln8$^{R24G}$ KI mice (D) while the littermate controls remain largely unaffected (C). A similar pattern was evident in the Cln8$^{R24G}$ male KI mice. 4 months: WT $n$ = 3, KI $n$ = 3; 9 months: WT $n$ = 4, KI $n$ = 4. Scale bars: Overview, 2.5 mm; hippocampus, thalamic nuclei and cortex, 100 µm.

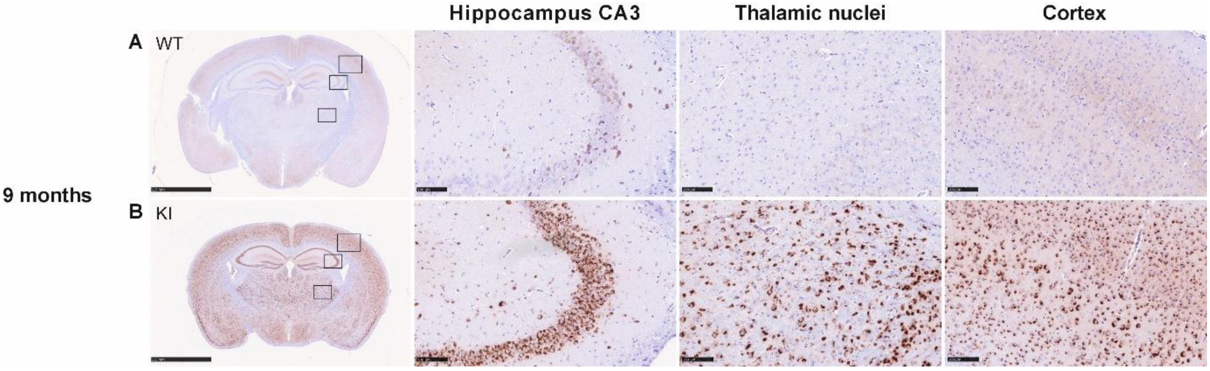


**Supplementary Figure 4. Immunohistochemical SCMAS staining of the hippocampal region of Cln8$^{R24G}$ B-line mice to visualise the accumulation of storage material.** The accumulation was prominent in the hippocampus, VPN/VPL thalamic nuclei and the cortex of the Cln8$^{R24G}$ KI mice (B) but not of the littermate controls (A) at 9 months of age (WT $n$ = 2, KI $n$ = 2). Scale bars: Overview, 2.5 mm; hippocampus, thalamic nuclei and cortex, 100 µm.

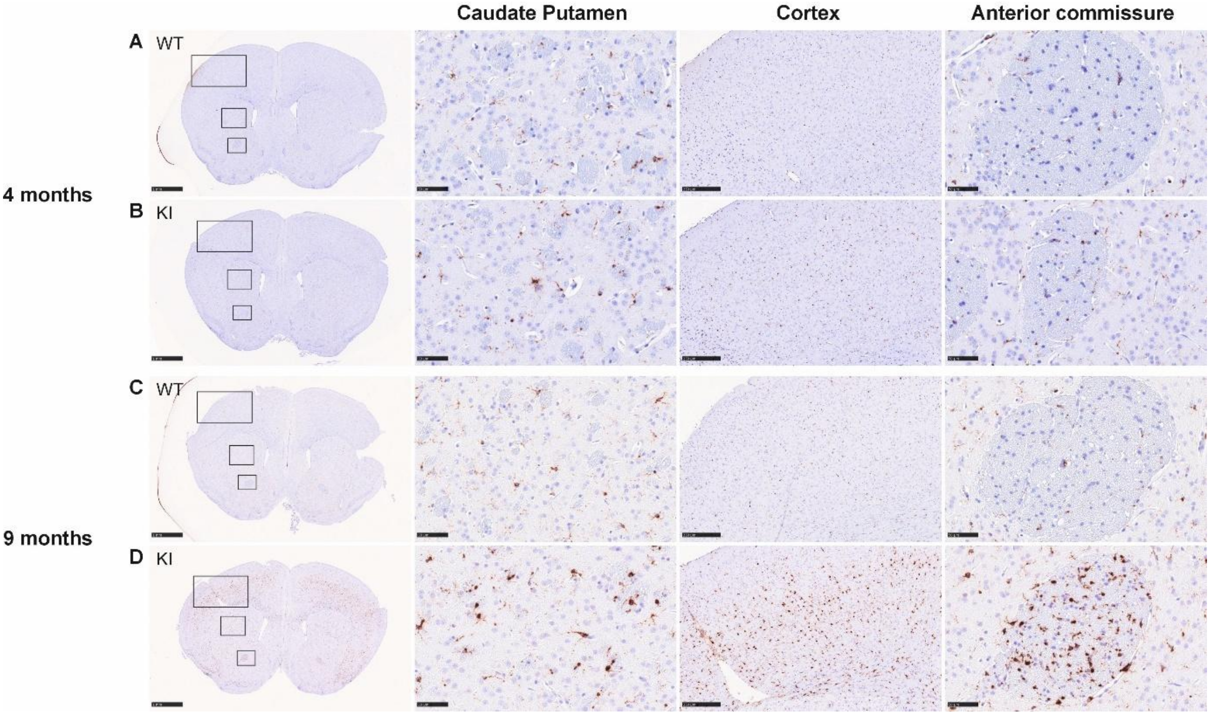


**Supplementary Figure 5. Immunohistochemical staining of the striatum region of Cln8$^{R24G}$ A-line males with marker for microglia (CD68)**. No clear differences in the number of CD68 positive (CD68$^{+}$) cells was evident in 4-month-old Cln8$^{R24G}$ KI mice in the striatum (A, B). However, a clear increase in CD68$^{+}$cells was observed in the motor cortex and the anterior commissure of Cln8$^{R24G}$ KI mice at 9 months of age (D) compared to the littermate controls (C). Only a slight increase in CD68$^{+}$ cells was observed in the caudate putamen of Cln8$^{R24G}$ KI mice (C, D). 4 months: WT *n* = 2, KI *n* = 2; 9 months: WT *n* = 3, KI *n* = 3. Scale bars: Overview, 1 mm; caudate putamen and anterior commissure, 50 µm, cortex, 250 µm.

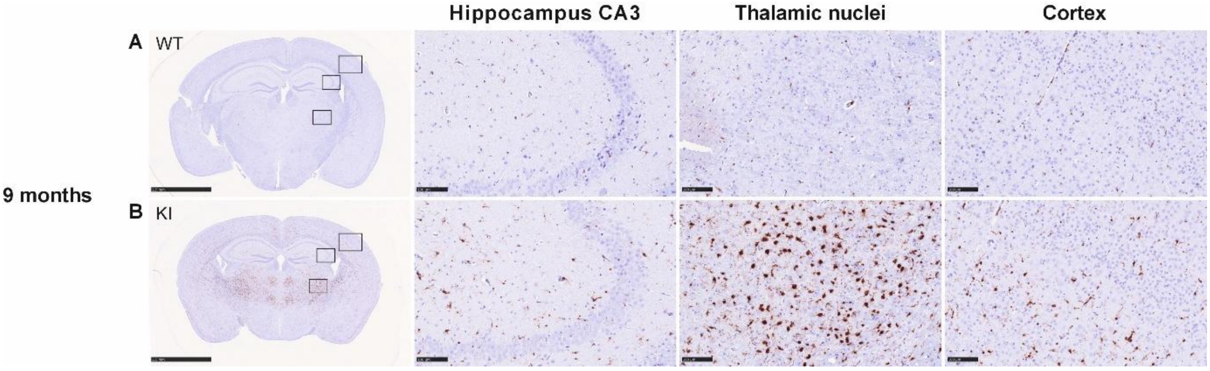


**Supplementary Figure 6**. **Immunohistochemical staining of hippocampal region of the Cln8$^{R24G}$ B-line males with marker for microglia (CD68)**. An increase in the number of CD68 -positive (CD68$^{+}$) cells was observed in the thalamic VP nuclei of the Cln8$^{R24G}$ KI mice (B) at 9 months of age compared to the littermate controls (A). An increase in CD68$^{+}$ cells was also noted in the somatosensory cortex of the Cln8$^{R24G}$ KI mice, though less prominently than in the thalamic nuclei (B). No difference in the number of CD68$^{+}$ cells was detected in the hippocampus of the Cln8$^{R24G}$ KI mice and the littermate controls (WT *n* = 2, KI *n* = 2). Scale bars: Overview, 2.5 mm; hippocampus, thalamic nuclei and cortex, 100 µm.

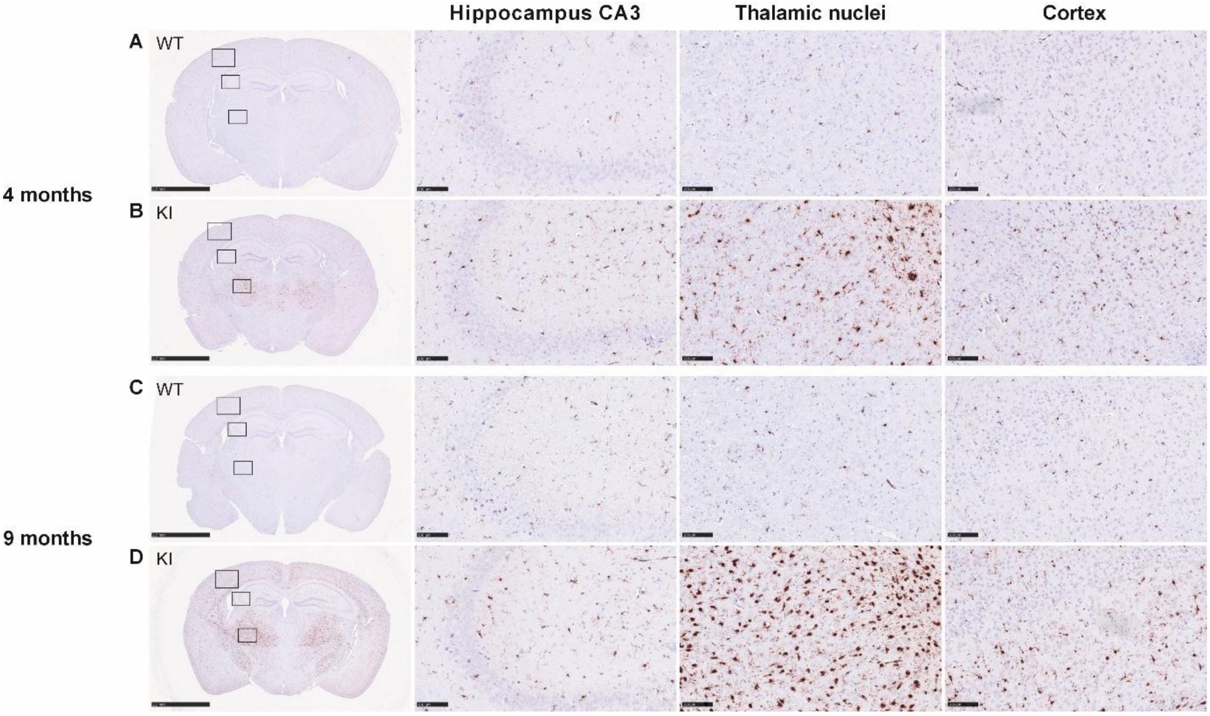


**Supplementary Figure 7. Immunohistochemical staining of hippocampal region of the Cln8$^{R24G}$ A-line females with marker for microglia (CD68).** A clear increase in the number of CD68 -positive (CD68$^{+}$) cells was evident in the thalamic VP nuclei of female Cln8$^{R24G}$ KI mice at 4 months and 9 months of age (as also observed in the male Cln8$^{R24G}$ KI mice) (B, D) compared to the littermate controls (A, C). The number of CD68$^{+}$ cells increases markedly in this region with age (B, D). An increase in CD68$^{+}$ cells was also noted in the somatosensory cortex of Cln8$^{R24G}$ KI mice at both ages though less prominent than in the thalamic nuclei (B, D). No clear difference in the number of CD68$^{+}$ cells was evident in the hippocampi of Cln8$^{R24G}$ KI mice (B, D) and the littermate controls (A, C) (4 months: WT *n* = 3, KI *n* = 3; 9 months: WT *n* = 4, KI *n* = 4). Scale bars: Overview, 2.5mm; hippocampus, thalamic nuclei and cortex, 100 µm.

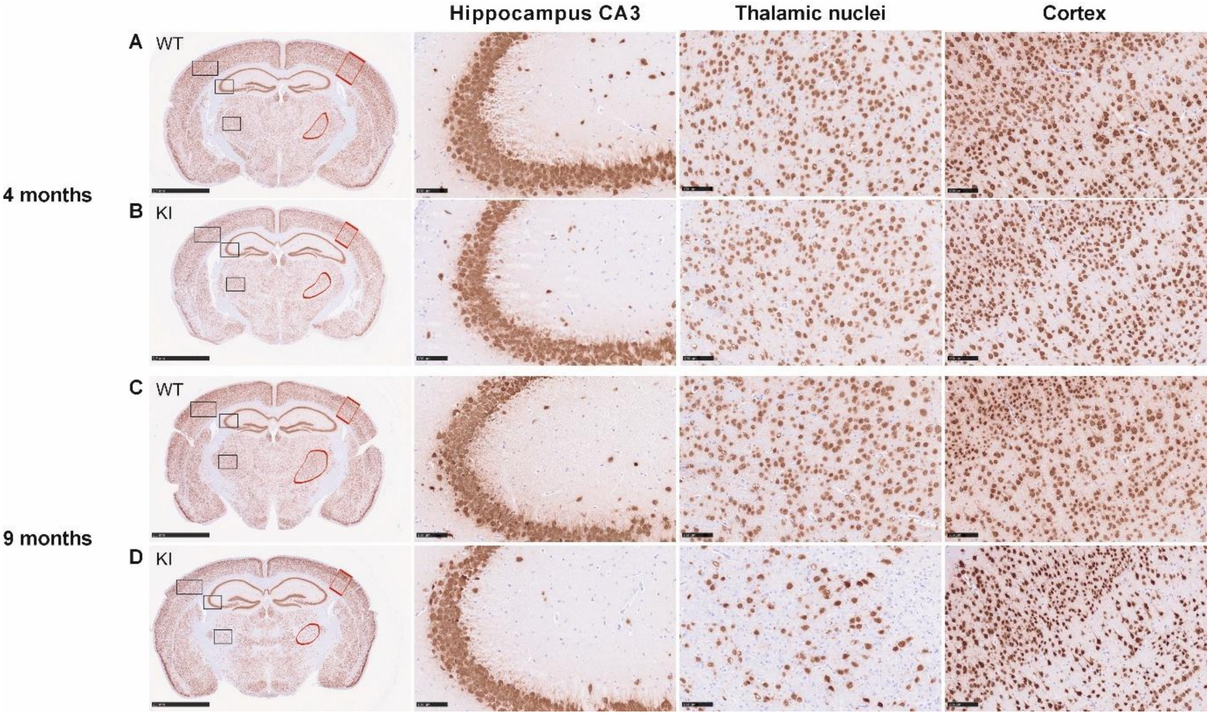


**Supplementary Figure 8. Immunostaining of hippocampal region of the Cln8$^{R24G}$ A-line females using an antibody against the neuronal marker NeuN.** Staining showed significantly decreased neuronal number in the thalamic nuclei of Cln8$^{R24G}$ female KI mice (D) at 9 months of age compared to the littermate controls (C). No clear differences in the neuronal number were seen in the hippocampus and cortex of Cln8$^{R24G}$ KI mice (B, D) compared to the littermate controls (A, C) at both age points. The black boxes indicate the area from where the close ups are taken and the red boxes indicate the areas used for the NeuN quantification.

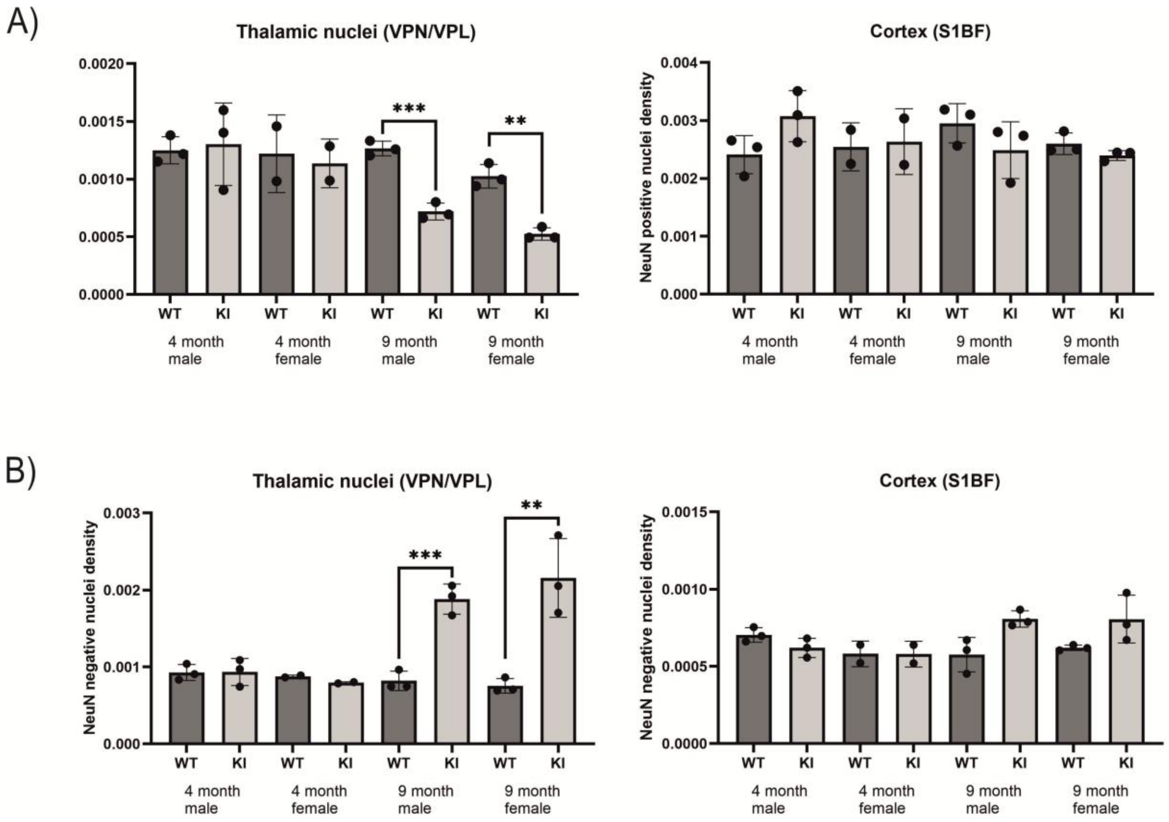


**Supplementary Figure 9. NeuN quantification in the left brain hemisphere of Cln8$^{R24G}$ KI A line mice and littermate controls.** A) A significant decrease in the density of the NeuN - positive nuclei (NeuN$^{+}$) was observed in the thalamic nuclei (VPN/VPL) of the Cln8$^{R24G}$ KI mice compared with the littermate controls at 9 months of age, whereas no differences were evident at 4 months. No differences in NeuN$^{+}$ nuclei density were detected in the cortex of the Cln8$^{R24G}$ KI mice at either age. B) A significant increase in the density of NeuN -negative nuclei (NeuN$^{-}$) was observed in the thalamic nuclei (VPN/VPL) of the Cln8$^{R24G}$ KI mice at 9 months. In the somatosensory cortex (S1BF), a non-significant increase in NeuN$^{-}$ nuclei density was evident at 9 months, but not at 4 months. *$P$ <0.05, **$P$ < 0.01, ***$P$ < 0.001
